# A murine model of *Shigella sonnei* intestinal colonization

**DOI:** 10.64898/2026.05.28.728598

**Authors:** Lulu Liu, Sarah E. Woodward, David Baker, Zhe Zhao, Suthida Chenklin, Emma Slack, Christoph M. Tang

## Abstract

*Shigella sonnei* is a leading cause of bacterial dysentery and a high priority WHO pathogen because of the spread of multidrug resistant strains. Understanding microbiome-*Shigella*-host interactions during colonization of the gastrointestinal tract, and the development of vaccines have been hampered by the lack of small animal models of shigellosis. Here, we developed a murine model of intestinal colonization with *S. sonnei*. Pre-treatment of mice with antibiotics disturbed the intestinal microbiome and rendered mice susceptible to high level, gastrointestinal colonization with *S. sonnei* for over one week. Infection with *S. sonnei* CS14 harbouring a stable virulence plasmid induced an initial inflammatory response in wild type mice, with weight loss and elevated levels of fecal lipocalin 2; the *S. sonnei* Type III Secretion System was responsible for this inflammatory response. Expression of O-antigen and Group IV capsule by *S. sonnei* promoted sustained intestinal colonization, with infected mice developing mucosal and systemic antibody responses predominantly directed at these glycans. Finally, infection with *S. sonnei* induced a degree of protection against subsequent re-challenge. Overall, this murine model successfully mimics aspects of *S. sonnei* colonization and should be helpful in understanding how *S. sonnei* successfully survives within the gastrointestinal tract and competes with the microbiota as well as the evaluation of vaccine candidates.

## INTRODUCTION

Shigellosis is an important public health problem caused by members of *Shigella* spp. which are responsible for more than 267 million cases of diarrhea globally every year.^1^ There are an estimated 210,000 deaths each year resulting from infection with *Shigella* spp.^2^ Of the four species, *Shigella flexneri* and *Shigella sonnei* account for over 90% of all cases of shigellosis.^3^ *Shigella* infection is most prevalent in low- and middle- income countries (LMICs) where *S. flexneri* is endemic and primarily acquired by ingestion of contaminated food or water. In wealthier countries, *S. sonnei* predominates, with the frequency of infection caused by this species increasing in several countries as sanitation improves.^3^ Of significant concern, multidrug resistant (MDR) strains of *S. sonnei* threaten to render current frontline antimicrobials ineffective. Reports of MDR *Shigella* have increased in Europe over the past decade, while extensively drug resistant (XDR) *S. sonnei* has emerged in the USA. As a result, *S. sonnei* has been designated as a priority pathogen for the development of novel interventions by the World Health Organization (WHO) and Centers for Disease Control.^4–6^

*Shigella* spp. evolved from *Escherichia coli* following the acquisition of a large (∼210 kb) virulence plasmid, pINV.^7^ The plasmid encodes a Type III Secretion System (T3SS) on a 30 kb pathogenicity island (PAI) which is essential host cell invasion and disease.^8^ Unlike other species of *Shigella*, *S. sonnei* pINV also contains the O-antigen biosynthesis operon, required for the synthesis of the O-antigen (O-Ag) of lipopolysaccharide (LPS). *S. sonnei* O-Ag consists of repeats of the disaccharide, N-acetyl-L-altrosaminuronic acid (LAltNAcA)/2-acetamido-4-amino-2,4,6-trideoxy-D-galactose (4-N-D-FucNAc), and is a major target for vaccine development.^9–11^ *S. sonnei* also expresses a group IV capsule (G4C), which consists of the same glycan linked to the outer membrane by an unknown mechanism; the G4C can protect the bacterium from complement-mediated killing and antibacterials found in the gastrointestinal tract such as bile acids and antimicrobial peptides.^12–14^

Understanding *S. sonnei* pathogenesis and the development of novel therapeutics and vaccines have been hampered by the limited availability of small animal models of infection.^15^ Several models have been to study *S. flexneri* infection. The Sereny test, keratoconjunctivitis in guinea pigs, has been used to assess the virulence of strains,^16^ similar to the mouse intrapulmonary challenge model;^17^ furthermore, intrarectal challenge of guinea pigs can elicit acute proctocolitis.^18^ A murine model has also been developed using antibiotic-pretreated, inflammasome-deficient mice, but longer term infection (of more than 2-3 days) has not been reported.^19,20^ Encouragingly, antibiotic treatment combined with zinc deficiency of wild-type mice offers a model with sustained intestinal colonization with *S. flexneri*.^21^

For *S. sonnei*, models have been developed with zebrafish larvae with bacteria either added to the water or injected systemically.^19,22^ Additionally, wild-type mice pretreated with streptomycin have been used to study the effect of *S. sonnei* Type VI Secretion System (T6SS) during intestinal colonization lasting one or two days.^23^ However, there are currently no available murine models to assess the contribution of bacterial factors to *S. sonnei* colonization and evaluate vaccine candidates over longer timescales.

The first step in pathogenesis of *S. sonnei* infection is the successful colonization of the gastrointestinal (GI) tract.24-26 Within the GI tract, *S. sonnei* encounters and must compete with the resident microbiota, an important aspect of host defensive barrier that can prevents invasion by a pathogen through multiple mechanisms.^27–30^ The microbiome can block pathogens by direct antagonism through contact-dependent or - independent mechanisms, such as T6SS or colicins, respectively.^31,32^ Additionally, the microbiome can compete with newly acquired species for environmental niches and essential nutrients including iron, carbon and nitrogen sources, and electron acceptors.^33–35^ As a result of significant competition from the microbiome, as well as differences in innate immune and epithelial biology, mice are generally resistant to infection with *Shigella* spp.^20,36,37^

Here, we developed a murine model of *S. sonnei* long-term colonization in wild-type 129S6/SvEv mice which are commonly used for studies of inflammatory bowel disease.^38–40^ Mice were given antibiotics in their drinking water for three days, then infected with *S. sonnei* by oral gavage. We took advantage of a strain that contains a stable version of pINV, so that infecting bacteria consistently expressed their T3SS and O-Ag. High-level colonization of the gastrointestinal tract with this strain was sustained, lasting over 7 days, with an initial, transient, T3SS-dependent local inflammatory response to infection. Without antibiotic pre-treatment to reduce colonization resistance, colonization only lasted a few days. Analysis of mutants in competitive infection with the wild-type strain demonstrated the importance of *S. sonnei* O-Ag and the G4Cduring intestinal colonization. We show that, similar to human infection, mice infected with *S. sonnei* develop local and systemic antibody responses predominantly directed towards the O-antigen and are partially protected against rechallenge. This model should be valuable for identifying bacterial factors that contribute to *S. sonnei* colonization, and evaluating vaccines against this important human pathogen.

## RESULTS

### Alteration of the microbiome allows sustained intestinal colonization with *S. sonnei*

To develop a murine model of *S. sonnei* colonization, we exploited the clinical isolate *S. sonnei* CS14, which is naturally resistant to several antibiotics.^41–44^ Additionally, while most *S. sonnei* strains rapidly use their pINV plasmid on culture or outside of a host, a single nucleotide polymorphism (SNP) in the *vapBC* toxin-antitoxin (TA) system on pINV, *S. sonnei* CS14 maintains the virulence plasmid at high levels.^45^ We used *S. sonnei* CS14 to infect 6 -10-week-old 129S6/SvEv mice.

Initially, we examined the ability of *S. sonnei* CS14 to colonize the gastrointestinal tract of mice that had not been given antibiotics. Following oral gavage with 2×10^8^ CFU of *S. sonnei* on day 0, colonization was assessed by plating feces on media containing carbenicillin (to select for *S. sonnei* CS14) and Congo red (CR); colonies of *S. sonnei* expressing a T3SS appear red on plates containing CR (**Figure 1A**).^46^ Without antibiotic pre-treatment, mice were colonized with *S. sonnei* at a level of ∼ 10^5^ CFU/g feces for a few days post-challenge (**Figure 1B**); from the second day after challenge onwards, the level of colonization decreased, and by day 7, most mice cleared *S. sonnei* infection.

**Figure 1.**
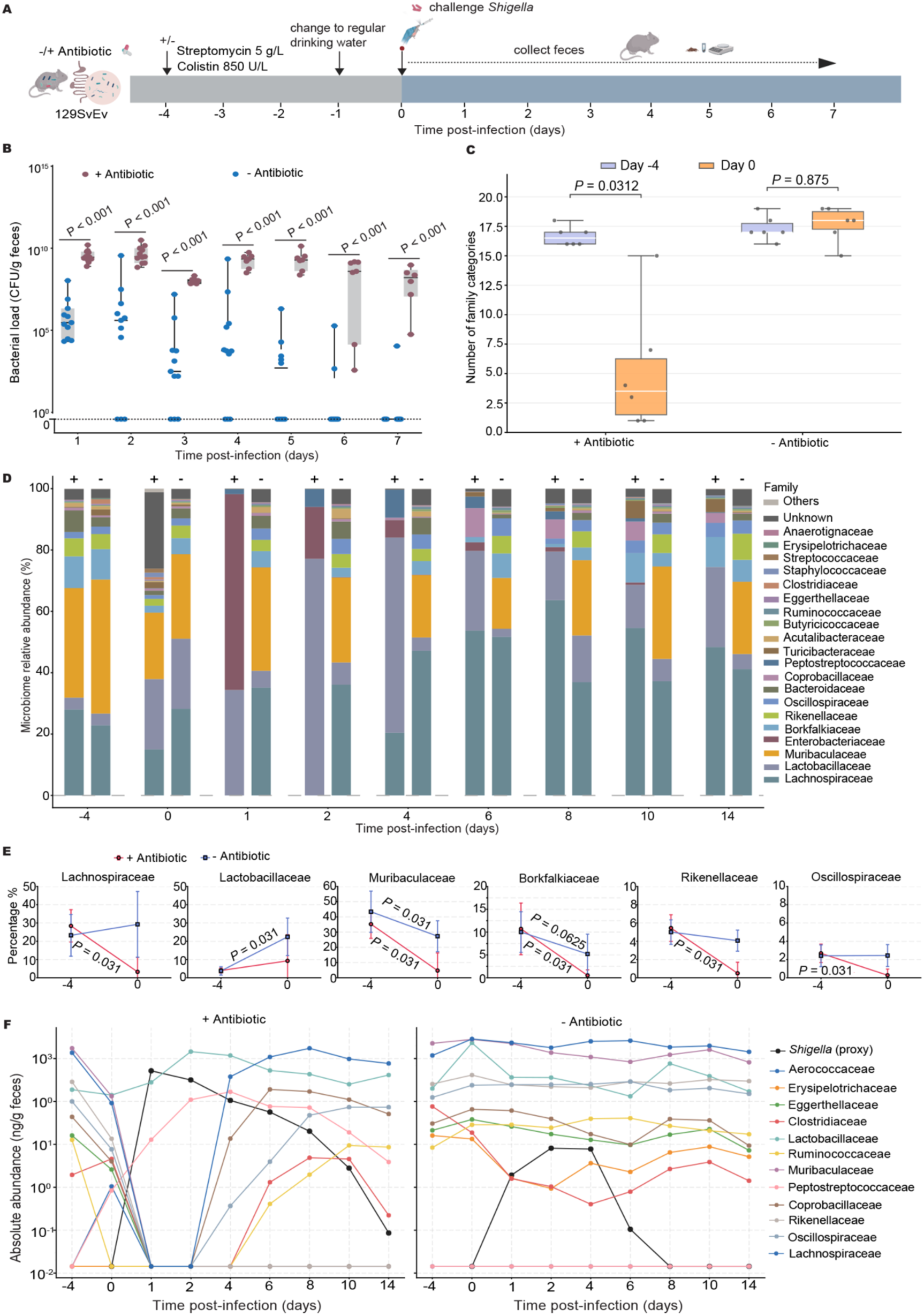
Development of a murine model for *S. sonnei* colonization. **A)** Mice received antibiotics in their drinking water on days -4 to -1, then were infected by oral gavage with 2 x10^8^ CFU *S. sonnei* CS14 on day 0. Following challenge, feces were collected daily, and **B)** *S. sonnei* carriage measured as CFU/per gram of feces in mice. Whiskers represent min to max. Dots represent individual mice; *n* = 11 per group; Mann-Whitney U). **C)** Number of family-level category between Day -4 and Day 0 in antibiotic-pretreated versus untreated groups. Each point is one mouse. *P* values are paired Wilcoxon signed-rank tests matching day -4 and day 0 by each mouse. **D)** Changes in the microbiota family relative abundance on days -4 to 14 following oral gavage with *S. sonnei* on day 0 with (+) or without (-) antibiotic pre-treatment. **E)** Relative percentage change of the top abundance families from D-4 to D0 with or without antibiotics treatment. Points show mean of family abundance of replicates; err bar, SD; *P* values are paired Wilcoxon signed-rank tests with each group. **F)** Absolute abundance (normalised with the weight of feces) of the main microbiota family changes from day -4 to day 14 of the groups with (+) or without (-) antibiotic pre-treatment align with the amount of *Shigella* species.

Pre-treatment with antibiotics can alter the gut microbiome and reduce its ability to provide colonization resistance against *S. sonnei.* To test this hypothesis, mice were given streptomycin (5 g/L) and colistin (850 U/L) in their drinking water from days -4 to -1. These antibiotics were selected as streptomycin has been used to impair colonization resistance against *Salmonella* and colistin is highly effective against a broad range of Gram negative bacteria.^47^ Following administration of these antibiotics, up to 10^9^ CFU of *S. sonnei*/g of feces were recovered consistently from mice given 2× 10^8^ CFU of *S. sonnei* on day 0 with high level colonization persisting for 7 days post challenge (**Figure 1B**).

To identify bacterial families affected by the administration of antibiotics, we performed metagenomic profiling of the fecal microbiome during antibiotic pre-treatment and subsequent *S. sonnei* challenge. In mice receiving antibiotics, there was a significant reduction in the number of detectable bacterial families from day -4 to day 0 (*P* = 0.031); in the absence of antibiotic treatment, the number of bacterial families remained stable over the same period as expected (*P* = 0.875, **Figure 1C**). We next profiled the relative and absolute abundances of bacterial families from day -4 to 14 (**Figure 1D, E**). Antibiotic treatment markedly disrupted the baseline microbiome composition between day -4 and day 0, resulting in a pronounced shift in the community structure (**Figure 1D**). Notably, the relative abundance of several dominant families declined significantly following antibiotic exposure, including *Lachnospiraceae* (28.4% to 3.3%, *P* = 0.031), *Muribaculaceae* (35.2% to 4.7%, *P* = 0.031), *Borkfalkiaceae* (10.7% to 0.5%, *P* = 0.031), *Rikenellaceae* (5.4% to 0.49%, *P* = 0.031), and *Oscillospiraceae* (2.68% to 0.27%, *P* = 0.031) (**Figure 1E**). Antibiotic treatment also led to a marked reduction in the absolute abundance of most bacterial families over the same period, with the notable exception of *Peptostreptococcaceae*, which underwent a pronounced expansion (**Figure 1F, TableS1**).

Following inoculation of antibiotic-treated mice with *S. sonnei*, there was a marked expansion in *Enterobacteriaceae*, which accounted for 80% of the fecal microbiota by day 1, consistent with successful colonization by *S. sonnei*. Most other bacterial families were virtually undetectable in both relative and absolute abundance at this time, except *Lactobacillaceae* which remained relatively stable throughout experiments. From day 2 onward, both the relative and absolute abundance of *Enterobacteriaceae* declined steadily, with recovery of the population of several commensal families, including *Lachnospiraceae*, *Coprobacillaceae*, and *Oscillospiraceae*. Furthermore, several bacterial families, including *Rikenellaceae* and *Bacteroidaceae*, failed to recover following antibiotic treatment (**Figure 1D, F**). In contrast, in mice that did not receive antibiotics, *S. sonnei* colonization had a minimal impact on the overall composition of the microbiome. Only two dominant families, *Erysipelotrichaceae* and *Clostridiaceae*, exhibited a transient decrease from day 1 to day 4, coinciding with a temporary increase in *Enterobacteriaceae* abundance during this period (**Figure 1F**).

### Infection with virulent *S. sonnei* elicits a transient inflammatory response

A hallmark of clinical shigellosis is the development of acute intestinal inflammation resulting in diarrhoea and weight loss.^3,21^ This is dependent on the ability of *S. sonnei* to invade intestinal epithelial cells through the activity of its T3SS.^48^ To explore the development of disease following inoculation with *S. sonnei*, both infected or mock-infected mice were weighed daily. Mice infected with wild-type *S. sonnei* lost between 5 to 10% of their body weight by days 1 and 2 post-challenge (**Figure 2A**). Interestingly, despite the low levels of gut colonization, animals without antibiotic pre-treatment also showed significant weight-loss, independent of high-level gut colonization. Weight loss was not observed in mice gavaged with media alone (weight of mice +antibiotic/+infection *vs.* +antibiotic/-infection, *P* < 0.001 and *P* = 0.08 on days 1 and 2, respectively). These data indicate that challenge with wild-type *S. sonnei* induces weight loss in mice, regardless of whether they had been pre-treated with antibiotics (**Figure S1A**).

**Figure 2.**
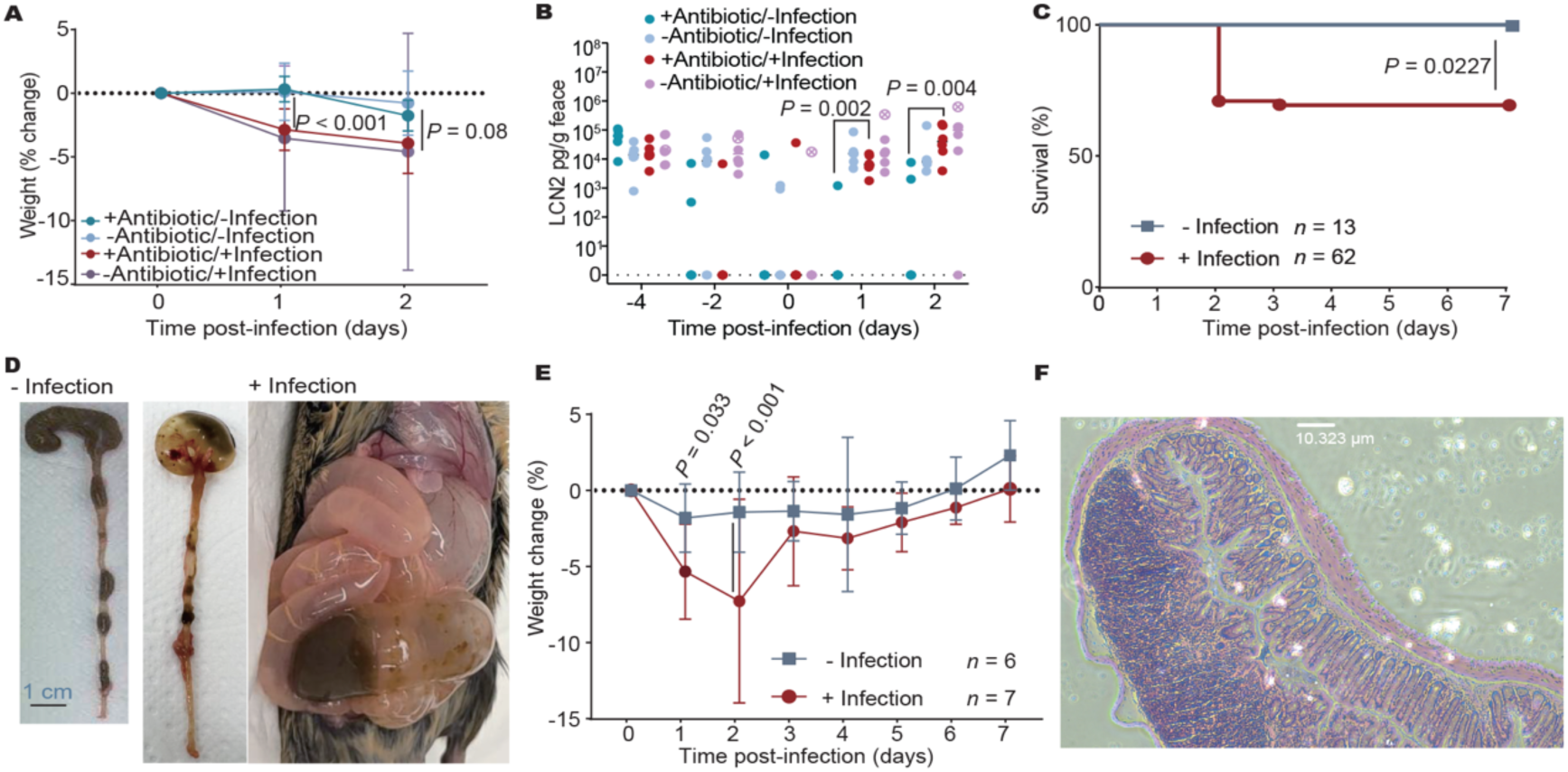
Infection with *S. sonnei* elicits a transient inflammatory response. **A)** Weight of mice given antibiotics (on days -4 to -1) or not, then gavaged with either 2 x10^8^ CFU of *S. sonnei* or media alone on day 0. **B)** Feces lipocalin 2 (LCN2) in mice challenged with *S. sonnei* or media alone with or without antibiotic pre-treatment. For A and B, *n* = 6/group; dots, show individual mice and the dot with cross showing significant higher lipocalin at humane endpoint; Mann-Whitney U. **C)** Percentage of mice with >15% loss of weight with (*n* = 62) or without (*n* = 13) *Shigella* infection. Log-rank test. **D)** Representative image illustrating intestinal abnormalities in morbid mice at humane endpoint. **E)** Weight of mice in a week with (*n* = 7) or without (*n* = 6) *Shigella* infection following antibiotic pretreatment. Dots, mean; error bars, SD; Two-way ANOVA, mixed model. **F)** Histology of infected mouse colon with prominent lymphoid.

To assess whether *S. sonnei* elicits intestinal inflammation, we measured levels of fecal Lipocalin-2 (LCN2), a marker of inflammation produced by neutrophils, epithelial cells, and hepatocytes.^49^ With or without antibiotic pre-treatment, infection of mice with *S. sonnei* led to significant increase in fecal LCN2 levels compared to uninfected mice on day 2 (*P* < 0.001, **Figure 2B**) (**Figure S1B**), indicating a local inflammatory response to challenge with *S. sonnei*.

A proportion mice infected with wild-type *S. sonnei* reached a humane endpoint (>15% weight loss) by day 2 (**Figure 2C**) and were euthanized in line with protocols; some of these mice displayed toxic megacolon upon necropsy (**Figure 2D**).^50^ Otherwise, the weight of surviving infected mice recovered by day 3 or 4, with fecal LCN2 levels returning to baseline (**Figure 2E**). We also examined histopathological changes in the colon of infected mice on day 2 when fecal LCN2 levels were at their peak; infected mice showed a trend for mild inflammation in the middle and distal colon with the mucosal layer distorted and a dense cellular infiltrate (**Figure 2F**), although the histology scores were not significantly different from those in uninfected mice (**Figure S2A**). To detect further evidence of inflammation, we measured cytokine responses in the distal colon. On day 2, mice were euthanized and tissue (∼ 0.5 cm in length) was isolated, washed then homogenized. However, there was no significant difference in the level of cytokines in infected and non-infected mice (**Figure S2B-D**), consistent with previous observations that *S. sonnei* induces less inflammation than other species such as *S. flexneri* and that host innate immunity successfully contains *Shigella* invasion in mice.^51^

Taken together, infection of mice with wild-type *S. sonnei* led to transient weight loss and increased fecal LCN-2 in some mice, indicating that bacterial challenge provokes a mild inflammatory response. However, in most cases the disease was mild and transient.

### The T3SS is required for the inflammatory response and sustained colonization in infected mice

To examine whether the inflammation and weight loss were linked to *S. sonnei* virulence, we examined the role of the pINV-encoded T3SS, which is necessary for bacterial entry into host cells and manipulation of immune cell signaling.^52,53^ We first compared the extent of *S. sonnei* colonization in mice challenged with wild-type *S. sonnei* (WT) or a T3SS-deficient mutant (T3SS^-^, through an IS*2*-mediated, 63,935 bp deletion, **Figure 3A**); both groups of mice were pre-treated with antibiotics (**Figure 3B**). Interestingly, there was no difference in the initial bacterial load of mice infected with the wild-type or T3SS^-^ strains (*P* = 0.165, 0.440, and 0.639 on days 1, 2, and 3, respectively). However, after day 3, the colonization level of the T3SS^-^ strain fell steadily and was significantly lower than the wild-type strain on days 4-7 (*P* < 0.05, **Figure 3C**). We also measured bacterial counts in homogenized tissues at different sites along the gastrointestinal tract on day 2 post-infection. The bacterial load recovered was similar in mice infected with wild-type and ΔT3SS strains (cecum, *P* = 0.589; feces, *P* = 0.485; proximal colon, *P* > 0.999; middle colon, *P* = 0.589; distal colon, *P* = 0.065, **Figure 3D**), indicating that most bacteria detected were not due to T3SS-dependent cell invasion. However, mice infected with wild-type *S. sonnei* lost significantly more weight than those infected with the T3SS^-^ mutant (*P* = 0.08 on day 1 and *P* = 0.0008 on day 2, **Figure 3E**), even though colonization levels were equivalent at day 2 (**Figure 3C, D**). Mice challenged with the T3SS mutant also displayed significantly lower levels of fecal LCN2 than mice receiving wild-type bacteria (**Figure 3F**, *P* < 0.001 on days 2 and 3). Taken together, these results indicate T3SS plays a role in the initial inflammatory response to infection and contributes to *Shigella* longer-term colonization of the gut lumen.

**Figure 3.**
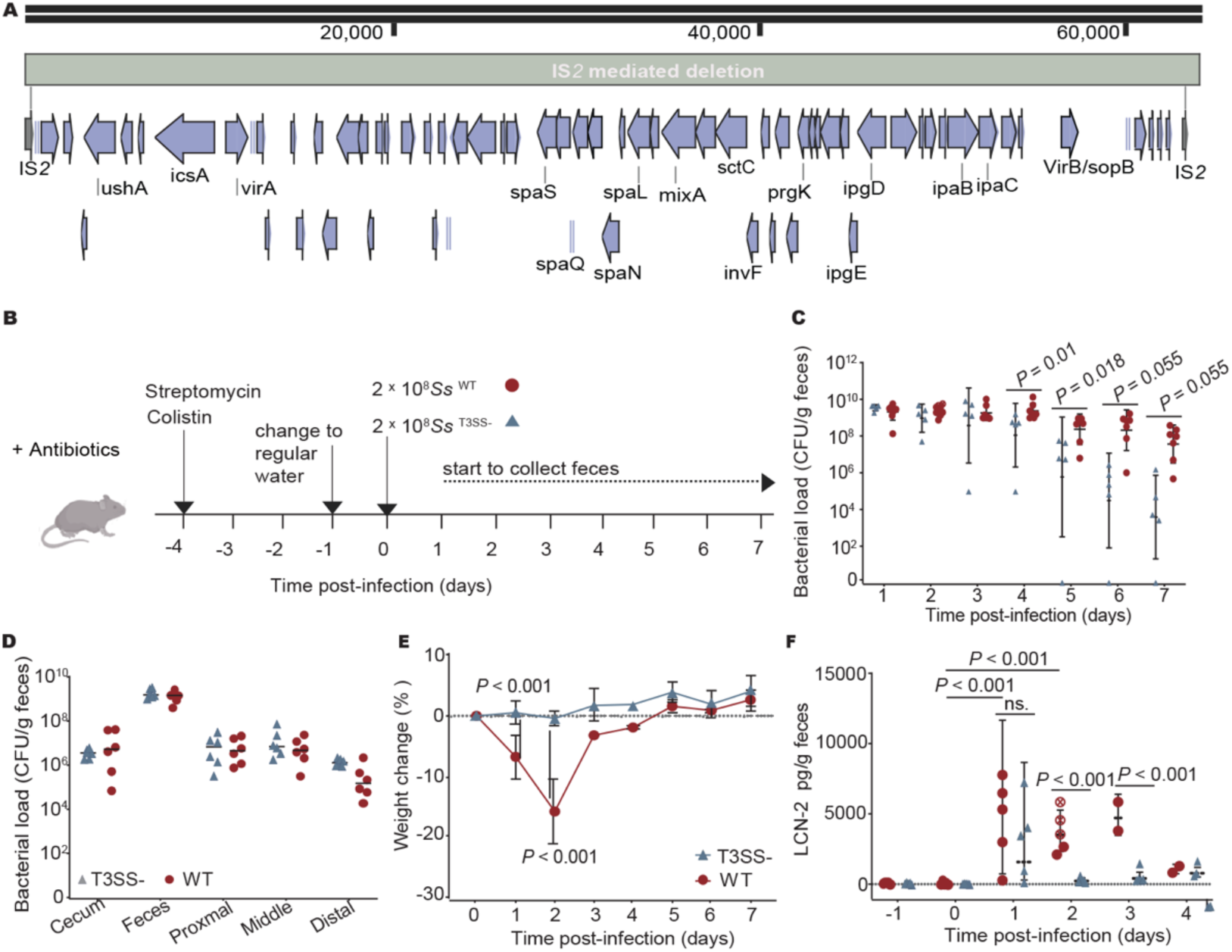
The *S. sonnei* T3SS contributes to the inflammatory response to colonization. **A)** Layout of IS*2* mediated T3SS system deletion. **B)** Intestinal colonization (CFU/gr of feces) of mice infected with wild-type (WT) or ΔT3SS mutant (ΔT3SS) *S. sonnei* CS14. *n* = 5-10; dots, individual mice, Mann-Whitney U. **C)** Bacterial load at sites in the large intestine; tissues were collected, washed and homogenized on day 2. *n* = 6; dots, individual mice. **D)** Bacterial load at sites in the large intestine; tissues were collected, washed and homogenized on day 2. *n* = 6; dots, individual mice. **E)** Weight of infected mice. *n* = 5; dots, mean; error bars, SD; Mann-Whitney U. **F)** Fecal LCN2 levels in mice infected with wild-type or ΔT3SS *S. sonnei*. dots with cross, individual mice euthanized in line with protocols. *n* = 5; horizontal lines, mean; error bars, SD; Two-way ANOVA, mixed model.

### Surface polysaccharides are essential for *S. sonnei* intestinal colonization

*S. sonnei* O-Ag and G4C both contain repeated LAltNAcA/4-N-D-FucNAc subunits (**Figure 4A**), which are produced by the O-Ag biosynthesis cluster on pINV;^54^ additional chromosomal genes are required to incorporate this glycan into the G4C.^55^ While loss of the G4C increases invasion of *S. sonnei* into epithelial cells, little is known about the role of the *S. sonnei* O-Ag and G4C during intestinal colonization. Therefore, we constructed in-frame deletion of the chromosomal G4C operon, removing *gfcA*-*gfcB*-*gfcC*-*ymcA*-*yccZ*-*etp*-*etk* to generate *S. sonnei* ΔG4C which produces an O-Ag but not a G4C (**Figure 4B**). A spontaneous O-Ag-deficient mutant (through an IS*91*-mediated 18,475 bp deletion) was selected from its colony phenotype; the O-Ag mutant is not predicted to express a G4C or O-Ag (**Figure 4C**). As expected, there was no difference in silver staining of LPS extracted from the wild-type and ΔG4C strains. However, LPS from the ΔOAg mutant lacked the characteristic ladder pattern of repeating O-Ag subunits (**Figure 4D**).

**Figure 4.**
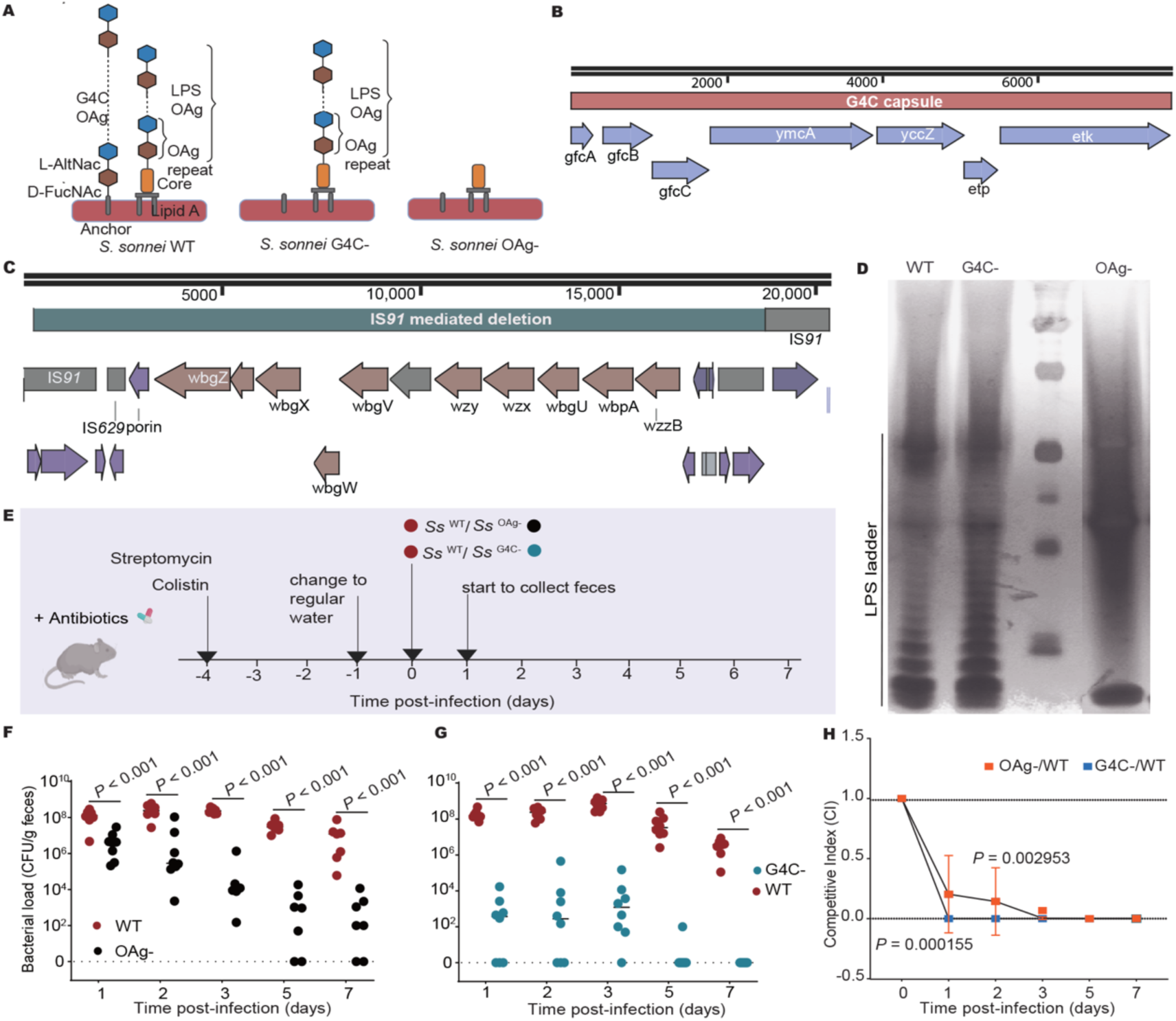
*Shigella* glycans influence intestinal colonization. **A)** Representation of the group IV capsule (G4C) and LPS O-antigen (OAg) of *S. sonnei* with relevant mutants indicated. **B)** Gene cluster of G4C synthesis pathway. **C)** IS*91* mediated O-antigen deletion. **D)** Profile of LPS from *S. sonnei* strains analysed by SDS-PAGE and silver staining. **E)** Mice were challenged with CS14WT and O-antigen mutant mix in 1:1 ratio of group 1 and CS14WT and G4C mutant mix in 1:1 ratio of group 2 respectively at day 0. **F, G)** Recovery of the ΔOAg and ΔG4C mutants compared to the wild type (WT). *n* = 8; dots, individual mice; Mann-Whitney U. **H)** Competitive index of the ΔOAg (orange) and ΔG4C (blue) mutants to WT. Dots, mean; error bars, SD; *n* = 8. Mann-Whitney U.

To establish whether surface gylcans expressed by *S. sonnei* contribute to intestinal colonization, mice were challenged with a ∼1:1 ratio of wild-type bacteria with either the ΔG4C or ΔOAg strain, and the ratio of mutant to wild-type in feces was determined over time, allowing calculation of the competitive index of the mutants (**Figure 4E**).^56^ Both the G4C and O-Ag mutants were rapidly outcompeted by the wild-type strain during colonization of the intestinal tract (ΔOAg *vs*. wild-type, *P* < 0.001; ΔG4C *vs*. wild-type, *P* < 0.001, **Figure 4F-G**), indicating that expression of surface glycans enhances bacterial fitness in this environment. Interestingly, the spontaneously arising ΔOAg strain, which lacks both the G4C as well as OAg, was outcompeted more slowly than the ΔG4C mutant, suggesting either that OAg production imposes a fitness cost or that another feature of the large genomic deletion in this strain has an associated fitness advantage (*P* < 0.001 on days 1 and 2, respectively, **Figure 4H**). ^57–60^

### Intestinal colonization with *S. sonnei* elicits antibody responses directed against bacterial OAg

Findings from epidemiological studies, the use of live attenuated vaccines, and controlled human infection models indicate that infection with *Shigella* spp. leads to serotype, O-Ag-specific protective immunity in humans.^61,62^ To examine whether mice colonized with *S. sonnei* develop adaptive immunity, local intestinal and systemic immune responses were measured at day 21 after challenge of mice with *S. sonnei* CS14; colonization with CS14 was cleared from day 7 to background levels by day 21 post-challenge (**Figure S3A, B**).

At 21 days post-challenge, levels of anti-*Shigella* antibodies were measured by flow cytometry of serum (IgG1, IgG2b, IgM) and IgA in the small intestine, using wild-type *S. sonnei* CS14 as the antibody binding target. Antibody levels were compared to those in control mice that had received antibiotics but not been challenged with *S. sonnei.* Compared to the uninfected group, significant intestinal and systemic antibody responses capable of binding wild-type *S. sonnei* CS14 were detected in mice infected with *S. sonnei* CS14 (IgA, *P* = 0.008; IgG1, *P* = 0.008, IgG2b, *P* = 0.008, **Figure 5A-D**).

**Figure 5.**
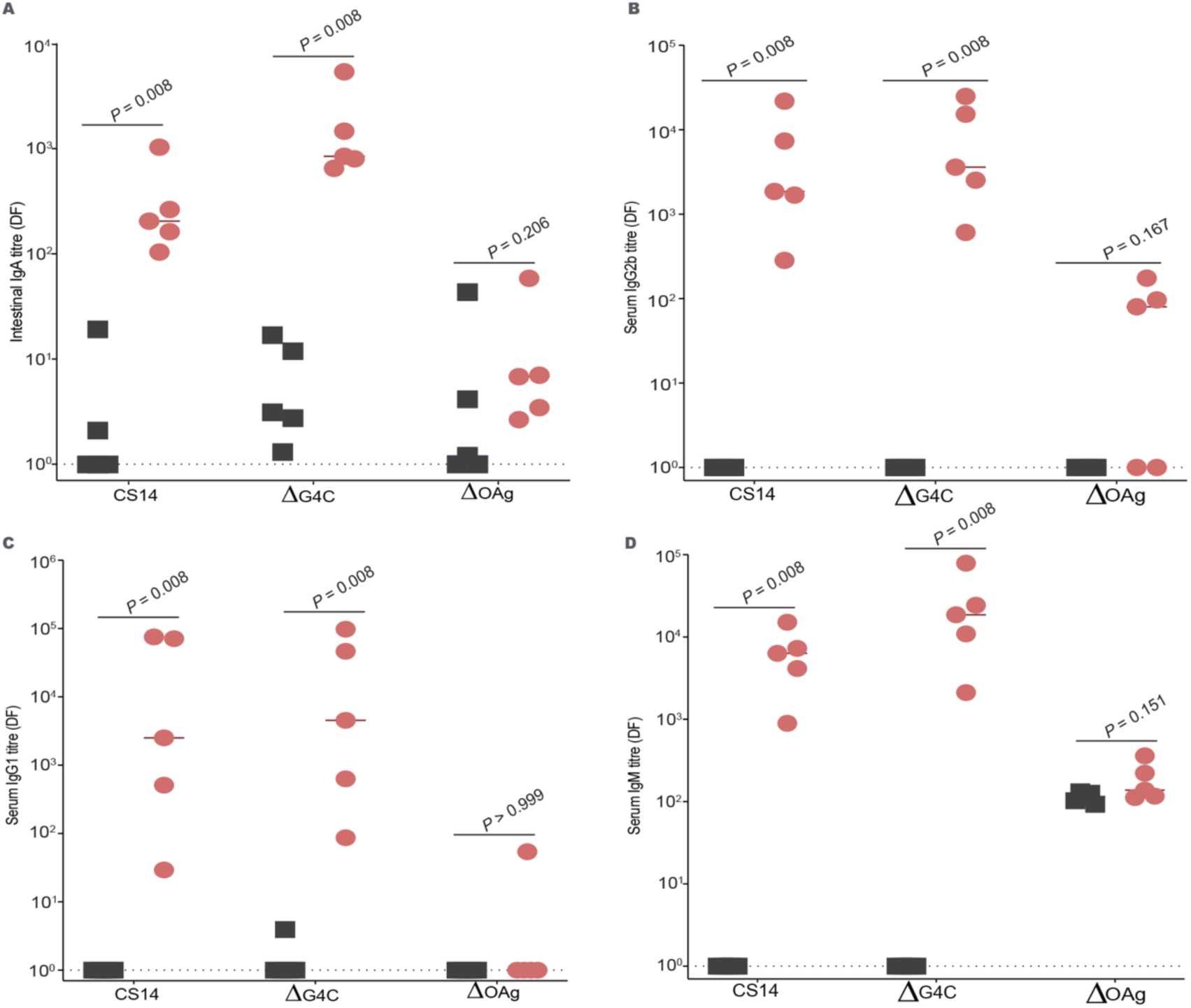
*S. sonnei* colonization elicits systemic and local immune responses against OAg. **A-D)** Antibody responses titre at day 21 following infection with *S. sonnei* CS14. Binding of Igs to *S. sonnei* CS14, ΔG4C or ΔOAg detected by flow cytometry. Dots, samples from individual mouse; *n* = 5; Mann-Whitney U test.

To further understand the antigenic targets of the anti-*Shigella* responses, we evaluated the ability of serum and mucosal antibodies to bind the ΔG4C and ΔOAg mutants. Although mucosal IgA, serum IgG2b, IgG1 and IgM of CS14 infected mice had a significant binding to the ΔG4C strain (IgA, *P* = 0.008; IgG1, *P* = 0.008; IgG2b, *P* = 0.008), binding of these antibodies to the ΔOAg mutant was barely detectable (IgA, *P* = 0.206; IgG1, *P* > 0.999; IgG2b, *P* = 0.151) (**Figure 5A-D)**. This indicates that the O-antigen is the major antigen recognized by the host immune system during intestinal colonization in the murine model, similar to human shigellosis.

### Prior infection protects mice from re-challenge

As infection with *S. sonnei* CS14 generates O-Ag-specific antibody responses, we next examined whether intestinal colonization with *S. sonnei* is protective against subsequent re-challenge (**Figure 6A**). To test this, groups of mice were pre-treated with antibiotics and received either vehicle alone (group 1, G1, no infection/infection group) or *S. sonnei* CS14 (G2, infection/re-infection group) on day 0. Mice were then kept for 17 days, before being given antibiotics in their drinking water (on days 17-20) to eliminate any residual *S. sonnei* from mice in G2, and make them susceptible challenge with *S. sonnei* CS14 on day 21 (**Figure 6A**). Of note, mice in G2 displayed significantly less weight loss following re-challenge (on days 22, 23 and 24, *P* = 0.01) and significantly lower colonization levels with wild-type bacteria (days 23 and 24, *P* = 0.005 and 0.013, respectively) compared to naïve mice (**Figure 6B-C**). These data indicate that *S. sonnei* carriage leads to the development of a degree of protective immunity in the murine model that manifests both as less inflammation and reduced intestinal colonization.

**Figure 6.**
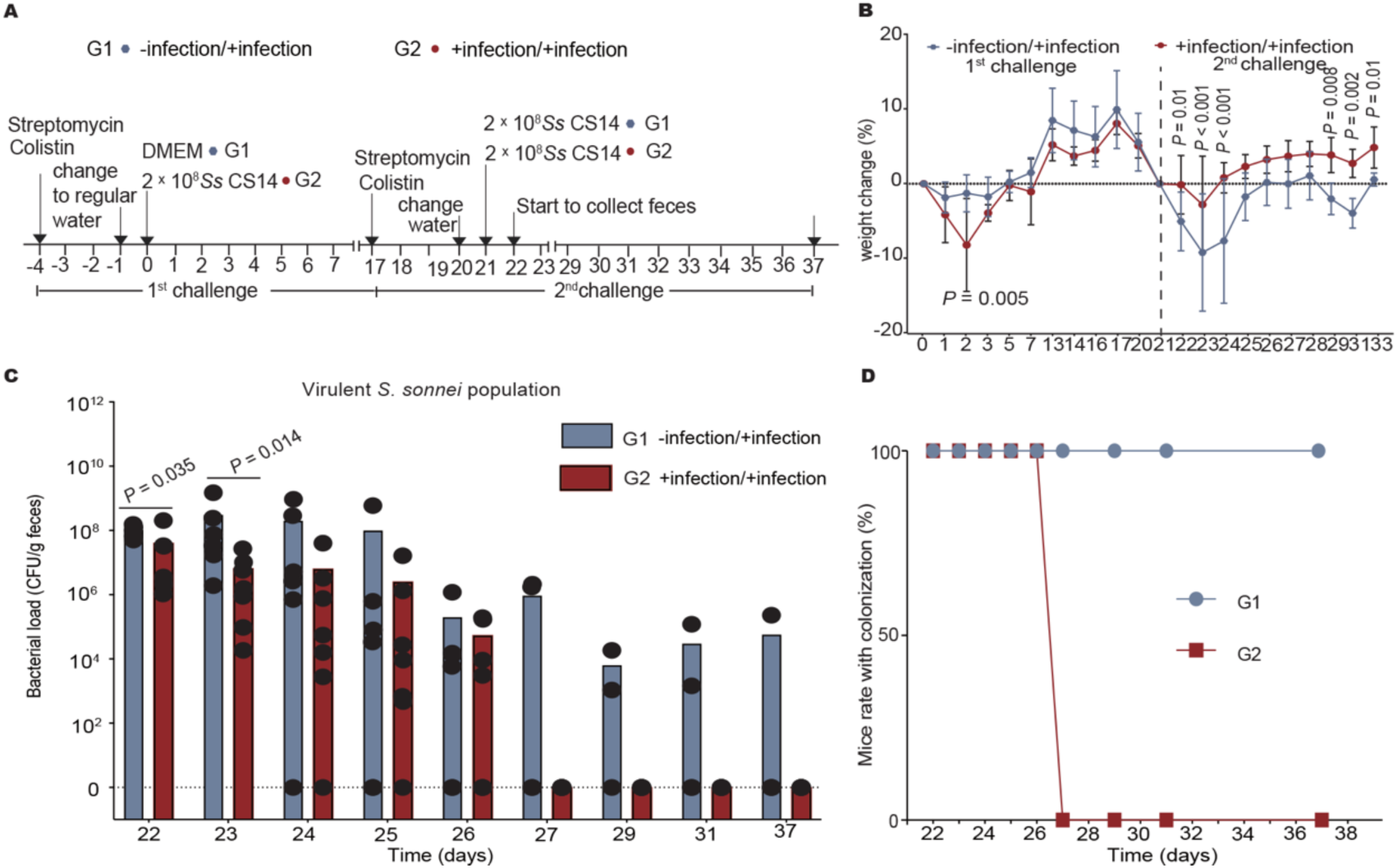
*S. sonnei* colonization confers protection against subsequent challenge. **A)** Mice received antibiotics on days -4 to -1, then mice in group 2 (G2) received oral gavage with *S. sonnei* on day 0, while those in group 1 (G1) were given DMEM only. From day 17 to 20, mice received antibiotics in their drinking water, then were challenged with *S. sonnei* CS14 by oral gavage on day 21. **B)** Weight of mice. Dots, mean; error bars, SD. *n* = 6 or 10. Two-way ANOVA, mixed model. **C)** Bacteria load of virulent *S. sonnei* following challenge of day 21. Dots, individual mice; *n* = 6 or 7. Mann-Whitney U. **D)** The percentage of mice colonized with virulent *S. sonnei* after challenge on day 21. The virulent colonies were confirmed by multiple PCR for pINV plasmid, genes of T3SS and O-antigen.

## DISCUSSION

*Shigella* is a leading cause of diarrhoea in LMICs and a high priority enteric pathogen, with *S. sonnei* outbreaks also observed in wealthy countries.^63^ With the rapid rise in AMR cases, there is an urgent need for better prevention and treatment of shigellosis. While several animal models have been developed *in vitro* and *in vivo* for *S. flexneri* to study pathogenesis and test vaccines, there is currently no murine model to study sustained *S. sonnei* colonization and assess vaccine candidates against this important cause of dysentery.

Here, we developed a murine model of *S. sonnei* intestinal colonization using wild-type 129S6/SvEv mice which have been commonly used for studying inflammatory bowel disease. Consistent with the commensal microbiota providing the first line of defence against shigellosis, we found that antibiotic pre-treatment of mice enabled *S. sonnei* CS14 to colonize the gastrointestinal tract at high levels for over 7 days. The antibiotic treatment regimen perturbed the resident microbiome, with metagenomic sequencing revealing that *Lactobacillaceae* was the first family to recover after cessation of antibiotic treatment and *Shigella* infection from day 1 to day 4. Initially, *Lachnospiraceae* was depleted, then recovered and became the dominant family from day 6 onwards, reaching levels equivalent to before antibiotic treatment (**Figure 1D, F**). *Lachnospiraceae* generate short chain fatty acids, which contribute to gut integrity, so its early re-emergence might curtail the inflammation seen following *S. sonnei* challenge.^64^

Following pre-treatment with antibiotics, oral gavage with *S. sonnei* CS14 induced mild yet detectable gastrointestinal inflammation with weight loss, peaking at two days post infection, similar to human disease. Interestingly, weight loss and increased fecal LCN2 were unaffected by antibiotic pre-treatment, indicating it was linked to bacterial challenge. The *Shigella* T3SS and its effectors encoded on pINV drive bacterial uptake by epithelial cells and survival within host cells.^48,65^ We found that the T3SS contributes to the early inflammation following challenge; this cannot be attributed to a reduced bacterial load, as during this initial period, mice infected with the T3SS mutant are colonized to similar levels as with the wild-type strain. Instead, the expression of the T3SS is associated with sustained colonization.

Infection with *S. sonnei* induced glycan-specific antibody responses, including IgA, IgM and IgG highlighting the potential for surface glycan vaccine-mediated immunity, which is being pursued by multiple approaches.^66,67^ Although *S. sonnei* CS14 consistently expresses a T3SS, we did not detect any responses against this essential virulence factor. We also observed that infection confers protection against *S. sonnei* re-challenge, presumably mediated by O-Ag specific immunity as seen in clinical infection.^61,67^

The model has some inherent limitations. First, inflammation following infection is mild with no obvious histopathological changes detected at day 2; we used immunocompetent mice and most rapidly recover following infection. Also, specific- pathogen-free mouse models have highly variable microbiota composition depending on the mouse vendor/breeding conditions. All experiments were carried out with one colony of mice, so antibiotic concentrations may need adapting for other background microbiota. Third, we have not explored the mechanisms of protection from virulent *S. sonnei* re-infection while it is tempting to speculate this is linked to the O-antigen-targeting antibody responses, given the short time between primary and secondary challenge, we cannot rule out a link to altered microbiome composition or innate immune priming.

Overall, this murine model successfully mimics aspects of *S. sonnei* infection disease, including transient inflammation and induction of adaptive immune responses directed at the O-Ag. Our findings highlight the importance of the commensal gut microbiota in preventing *S. sonnei* colonization. In the future, the model could be employed to identify Shigella colonisation factors and evaluate the ability of vaccines or drugs to prevent or eliminate gut colonization.

## MATERIALS AND METHODS

### Bacterial strains and growth conditions

All strains and plasmids used in this study are listed in **Table S2**. *S. sonnei* CS14::Carb^R^ and *S*. *sonnei* 53G::Kan^R^ were generated by inserting the selection markers (carbenicillin resistance, Carb^R^ or kanamycin resistance, Kan^R^) on chromosome between *gymA* and *aroG* (**Figure S5**) by conjugation with *E. coli* MFD pir Δ*hsdR* containing pCONJ derivatives;^68–70^ all primers are listed in **Table S3**. Bacteria were grown overnight on Tryptic soy broth (Merck, TSB) agar containing 0.1% CR (Sigma, CAS-No: 573-58-0) with 100 µg/mL carbenicillin at (Apollo Scientific Ltd, CAS-No: 4800-94-6) or kanamycin (Sigma, CAS-No:25389-94-0) as needed. Single colonies were grown in TSB liquid overnight at 37 °C with shaking at 180 r.p.m. Bacterial dilutions were prepared in phosphate buffered saline (PBS, Sigma, Cat. No. D8537-500ML).

### Murine model of shigellosis

Animal experiments were performed at the University of Oxford with 129S6SvEv mice in accordance with a Project License (PP5162604) approved by the Home Office of the UK. Specific pathogen-free 129mice, aged 6-10 weeks old, were bred in house. Mice were maintained on a 12-hour light-dark cycle. Female mice were randomly assigned to experimental and control groups. All animals were allowed free access to food and water throughout the duration of the experiment.

Antibiotics were added to the drinking water at 5 g/L streptomycin (Sigma, CAS: 3810-74-0) and 850 U/L colistin (Sigma, CAS: 1264-72-8). Antibiotic water was administered on days -4 to -1, after which all mice received standard, unmedicated water for the remainder of experiments.

Unless otherwise stated, wild-type *S. sonnei* was streaked from frozen stock onto TSA plates with 0.1% CR and incubated overnight at 37 °C. A single red, T3SS-positive colony was selected and inoculated into 5 mL TSB medium in a 30 mL universal tube, and incubated overnight at 37 °C with shaking at 180 r.p.m. The overnight culture was sub-cultured into fresh 5 mL TSB medium in a 30 mL universal tube and grown at 37 °C with shaking at 180 r.p.m. When the culture reached an OD₆₀₀ of ∼ 1.0, the suspension was diluted to 2 × 10⁸ CFU/100 µL in DMEM (Sigma, Cat. No. 5796) and 100 µL given to mice by oral gavage with a feeding tube (Instech, FTP-20-30-50). Competition Index (CI) was calculated as follows: the number of mutant/numbers of wild-type in samples divided by the number of mutants/numbers of wild-type in the inoculum.

### Bacterial load in feces or tissues

Fecal pellets were collected into pre-weighed 2 mL safe lock tubes containing 1 mL sterile PBS. Mice colon tissues (∼ 1 cm) were collected into pre-weighted 2 mL safe lock tubes containing 1 mL sterile PBS and tungsten bead. Samples were homogenized for 2 minutes at 25 Hz in a TissueLyser III (QIAGEN). Ten-fold serial dilutions were spotted in triplicate on TSA with 0.1% Congo red with appropriate antibiotics, then incubated in 37 °C and allowed to grow for 16 hours before enumeration of colony forming units (CFU). CFU were normalized by sample weight. Wild-type strains were counted by red and smooth colonies; the T3SS mutant colonies were determined with the white/pink smooth colonies and the OAg mutant colonies were counted with red rough colonies.

### Analysis of LPS expression

Bacteria were grown in 5 mL TSB overnight at 180 r.p.m in 37 °C incubator. Next day, bacteria were diluted 1:100 in 10 mL fresh TSB and grown at 180 r.p.m at 37 °C until the OD600 reached 1.0. Cells were harvested by centrifugation at 13,000 r.p.m for 2 min, and the pellet washed twice with PBS, then resuspended in 100 μL PBS. This was mixed with 100 μL sample buffer (6% SDS, 6% 2-mercaptoethanol, 10 mM DTT, 46% glycerol, 60mM Tris pH 8.0, 0.1% bromophenol blue) and heated to 98 °C for 10 min. Proteinase K (0.25% v/v, QIAGEN, cat. RP107B-5) was added and incubated at 37 °C overnight. Further proteinase K was added and the sample incubated at 55 °C for 3 hours, before visualization by SDS-PAGE (12% resolving gel and 4% stacking gel) and silver staining (Thermo Fisher, Pierce Sliver Stain Kit, cat. 24612).

### Detection of fecal LCN-2

Homogenized fecal pellets were homogenized for 2 minutes at 25 Hz in a TissueLyser III (QIAGEN). Then the samples were centrifuged at 13,000 *xg* for 2 min, then the supernatant analysed with the Mouse Lipocalin-2 ELISA Kit (NGAL) (ab119601) according to the manufacturer’s instructions. [how result was read and in what machine]

### Histopathological analysis of infected tissue

Colonic tissue was fixed in 4% paraformaldehyde (PFA) for > 24 hours then transferred into 70% ethanol before processing using a Tissue TeK VIP 6 Processor Sakura. Tissue samples were dehydrated through graded ethanol according to standard procedures. Samples were incubated in 70% ethanol for 1 h, then at 40 °C for 30 min, then transferred to 100% ethanol for 1 hour with an additional 30-minute incubation at 40 °C. The 100% ethanol step was repeated to ensure complete dehydration, and the tissue transferred to xylene (X/0250/21, Fisher) for 1 h then at 40 °C for 30 min (repeated three times), then paraffin (CellWax Plus (S), GCA-0400-00A, CellPath) for 1 h then 30 min at 63 °C (repeated four times) then left in paraffin at 63 °C overnight. Samples were finally embedded using TEC 5 Embedding centre (Sakura). Samples were treated in Xylene (Fisher, X/025021) for 3 min twice then treated in 100% ethanol twice, and strained with Haematoxylin Harris (Leica microsystems, 3801560E). Samples were treated with acid alcohol (1.4 L 100% ethanol with 600 mL dH2O and 20 mL 37% HCl) for 40 secs, washed, then incubated with Blueing Agent (600 mL dH2O with 1.6 mL ammonia (VWR, 87766.20)) for 1 min, and finally eosin Y (1%, Cell Path, RBC-0100-00A). Samples were dehydrated in 100% ethanol for 1 min, three times and xylene twice, before mounting on slides (mounting media in the G2 Cover slipper Machine Sakura). The samples were blinded and scored (Table S4).

### Microbiome analysis

Genomic DNA was extracted from fecal samples with the QIAamp PowerFecal Pro DNA Kit (Qiagen; cat. no. 51804) and analysed with a Nano Drop ND-1000 spectrophotometer (Thermo Fisher Scientific, Waltham, MA, USA). Libraries were prepared with Illumina DNA Prep Kit, quantified using Qubit 4.0 Fluorometer (Life Technologies, Carlsbad, CA, USA) and checked with an Agilent 5600 Analyzer (Agilent Technologies, Palo Alto, CA, USA). Sequencing libraries were multiplexed and loaded on the Illumina NovaSeq XPlus instrument according to manufacturer’s instructions. Samples were sequenced using a 2x150 Pair-End (PE) configuration; base calling was conducted by the NovaSeq Control Software v1.3 on the NovaSeq instrument. Raw sequence data (bcl files) were converted into fastq files and de-multiplexed using Illumina bcl2fastq program. One mismatch was allowed for index sequence identification.

The original raw data quality was assessed with Trimmomatic v0.36 software,^71^ removing adaptors, primers, and low-quality (Q < 20) data. The filtered data were subjected to quality assessment and basic statistical information using SeqKit v0.9.3 software. The relative abundance of known and uncharacterized microbial community members at the species level were analyzed statistically with mOTUs v3 tool,^72^ which utilizes ten single-copy phylogenetic marker genes for taxonomic analysis. These marker genes were extracted from over 25,000 reference genomes and 3,100 metagenomic samples and clustered into more than 7,700 MG-based operational taxonomic units (mOTUs) to represent different microbial species employing mOTUs v3.

### Detection of antibody responses by flow cytometry

Bacteria were grown overnight in LB media before diluting 1:3 in PBS 0.002% sodium azide (PBS-azide) to prevent internalization of bacterial surface antigens and pasteurized for 30 minutes at 60 °C. Cultures were prepared for flow cytometric analysis as described by Moor *et al*. (2016).^73^ Briefly, pasteurized bacteria were washed twice in PBS-azide (centrifugation at 7000 *xg* for 2 minutes) before dilution to 2.5x105 CFU per well in PBS-azide. Prepared bacteria were incubated for 30 minutes with murine intestinal lavage samples through eight 3-fold serial dilutions in PBS-azide. In the case of murine serum antibodies, samples were first diluted 1:100 in PBS-azide and heat-inactivated for 10 minutes at 56 °C prior to dilution and incubation with target bacteria. Samples were washed twice in PBS-azide to remove unbound antibody and resuspended in PBS-azide containing secondary BV421 rat anti-mouse IgA (1:50; BD Biosciences; cat. no. 743293; clone C10-1) for intestinal samples, or a combination of FITC rat anti-mouse IgG2b (1:100; BD Biosciences; cat. no. 553395; clone R12-3), PE rat anti-mouse IgG1 (1:100; BD Biosciences; cat. no. 550083; clone A85-1), and APC rat anti-mouse IgM (1:100; BD Biosciences; cat. no. 550676; clone II/41) for serum samples. Samples were incubated with secondary antibodies at 4 °C for 30 minutes to overnight before washing and resuspending in 4% paraformaldehyde pH 7.2 for acquisition of 10,000 events per sample on a CytoFLEX LX flow cytometer. Analysis was carried out in FlowJo 10.10.0, gating on bacteria before determination of median fluorescence intensity (MFI) on each fluorophore.

### Measurement of serum and colon tissue cytokine concentrations

Cytokine analysis was performed on both serum and colon tissues at endpoint on day 2 post-infection. Blood was collected into protein Lobind safe lock tubes (Eppendorf, Catalog No. 0030108094) and left at room temperature for 30 minutes then centrifuged at 2,000 *x g* for 10 minutes at 4 °C, and the serum collected. Murine colonic tissue (∼ 1 cm) was washed twice in PBS to remove fecal content and collected into pre-weighted 2 mL safe lock tubes containing 1 mL cOmplete EDTA-free protease inhibitor cocktail (Roche 04693159001). Buffer was prepared fresh on the day of sample collection. Samples were homogenized for 2 minutes at 25 Hz in a TissueLyser III (QIAGEN). Cytokines were measured using a Mouse Th1/Th2/Th17 Cytokine Bead Array (CBA) Kit (BD Biosciences; Cat. No. BDB560485), according to the manufacturer’s instructions followed by analysis using a CytoFLEX LX flow cytometer and analysis in FlowJo 10.10.0. Cytokine concentrations were normalized to tissue weight.

### RT-qPCR

Distal colon (∼0.5 cm) was collected into pre-weighted 2 mL safe lock tubes with 500 μL RNA*later* Solution (Cat. AM7020, Invitrogen), and RNA extracted using the FastPure Cell/Tissue Total RNA Isolation Kit V2 (Cat. RC112-01, Vazyme), according to the manufacturer’s instructions. Reverse transcription and real-time PCR was performed with SYBR Green (HiScript IV SuperMix for qPCR Cat: R4423-01 and Taq Pro Universal SYBR qPCR Master, Vazyme) according to the manufacturer’s instructions using the Roche LightCycler 480 System (Roche). Figure were generated directly with CT values of each sample.

### Statistical analyses

All statistic tests used are clarified in their respective figure legend. *P* values of less than 0.05 were considered statistically significant. Fold changes in mRNA levels, bacterial loads, competitive indices, and lipocalin concentrations underwent natural logarithmic transformation prior to statistical analysis, to account for log-normal error distributions generated by serial dilution.

### Data Availability

All data are available in the main text or supplemental information. Metagenomic sequencing raw data is available under BioProject: PRJNA1374196.

## Acknowledgements

We thank the all the staff in flow cytometry facility of Sir William Dunn School of Pathology, Animal Research Center and Histology lab from Kennedy Institute of Rheumatology, University of Oxford.

## Conflict of Interest Statement

The authors declare no competing interests.

## Notes

### Competing Interest Statement

The authors have declared no competing interest.

## REFERENCES

1. Badr, H.S., Colston, J.M., Nguyen, N.H., Chen, Y.T., Burnett, E., Ali, S.A., Rayamajhi, A., Satter, S.M., Van Trang, N., Eibach, D., et al. (2023). Spatiotemporal variation in risk of *Shigella* infection in childhood: a global risk mapping and prediction model using individual participant data. Lancet Glob Health. 11(3), e373–e384. 10.1016/S2214-109X(22)00549-6.

2. Khalil, I.A., Troeger, C., Blacker, B.F., Rao, P.C., Brown, A., Atherly, D.E., Brewer, T.G., Engmann, C.M., Houpt, E.R., Kang, G., et al. (2018). Morbidity and mortality due to shigella and enterotoxigenic *Escherichia coli* diarrhoea: the Global Burden of Disease Study 1990–2016. Lancet Infect Dis. 18(11), 1229–1240. 10.1016/S1473-3099(18)30475-4.

3. Scott, T.A., Baker, K.S., Trotter, C., Jenkins, C., Mostowy, S., Hawkey, J., Schmidt, H., Holt, K.E., Thomson, N.R., and Baker, S. (2024). *Shigella sonnei*: epidemiology, evolution, pathogenesis, resistance and host interactions. Nat Rev Microbiol. 23(5), 303–317. 10.1038/s41579-024-01126-x.

4. Mason, L.C.E., Greig, D.R., Cowley, L.A., Partridge, S.R., Martinez, E., Blackwell, G.A., Chong, C.E., De Silva, P.M., Bengtsson, R.J., Draper, J.L., et al. (2023). The evolution and international spread of extensively drug resistant *Shigella sonnei*. Nat Commun. 14(1), 1983. 10.1038/s41467-023-37672-w.

5. Bardsley, M., Jenkins, C., Mitchell, H.D., Mikhail, A.F.W., Baker, K.S., Foster, K., Hughes, G., and Dallman, T.J. (2020). Persistent Transmission of Shigellosis in England Is Associated with a Recently Emerged Multidrug-Resistant Strain of *Shigella* sonnei. J Clin Microbiol. 58(4):e01692–19. 10.1128/JCM.01692-19.

6. Caldera, J.R., Shaw, B., Uslan, D.Z., and Yang, S. (2024). Cluster of extensively drug-resistant *Shigella sonnei* carrying bla(CTX-M-15) in Los Angeles, California, 2023 to 2024. Am J Infect Control. 53(4), 524–526. 10.1016/j.ajic.2024.12.005.

7. The, H.C., Thanh, D.P., Holt, K.E., Thomson, N.R., and Baker, S. (2016). The genomic signatures of *Shigella* evolution, adaptation and geographical spread. Nat Rev Microbiol. 14(4), 235–250. 10.1038/nrmicro.2016.10.

8. Pilla, G., McVicker, G., and Tang, C.M. (2017). Genetic plasticity of the *Shigella* virulence plasmid is mediated by intra- and inter-molecular events between insertion sequences. PLoS Genet. 13(9), e1007014. 10.1371/journal.pgen.1007014.

9. Liu, B., Knirel, Y. A., Feng, L., Perepelov, A. V., Senchenkova, S. N., Wang, Q., Reeves, P. R., & Wang, L. (2008). Structure and genetics of *Shigella* O antigens. FEMS Microbiol Rev. 32(4), 627–653. 10.1111/j.1574-6976.2008.00114.x.

10. Shepherd, J.G., Wang, L., & Reeves, P. R. (2000). Comparison of O-antigen gene clusters of *Escherichia coli* (*Shigella*) *sonnei* and *Plesiomonas shigelloides* O17: *sonnei* gained its current plasmid-borne O-antigen genes from *P. shigelloides* in a recent event. Infect Immun. 68(10), 6056–6061. 10.1128/IAI.68.10.6056-6061.2000.

11. Rossi, O., Citiulo, F., Giannelli, C., Cappelletti, E., Gasperini, G., Mancini, F., Acquaviva, A., Raso, M.M., Sollai, L., Alfini, R., et al. (2023). A next-generation GMMA-based vaccine candidate to fight shigellosis. NPJ Vaccines 8(1), 130. 10.1038/s41541-023-00725-8.

12. Caboni, M., Pedron, T., Rossi, O., Goulding, D., Pickard, D., Citiulo, F., MacLennan, C.A., Dougan, G., Thomson, N.R., Saul, A., et al. (2015). An O antigen capsule modulates bacterial pathogenesis in *Shigella sonnei*. PLoS Pathog. 11(3), e1004749. 10.1371/journal.ppat.1004749.

13. Whitfield, C. (2006). Biosynthesis and assembly of capsular polysaccharides in *Escherichia coli*. Annu Rev Biochem. 75, 39–68. 10.1146/annurev.biochem.75.103004.142545.

14. Peleg, A., Shifrin, Y., Ilan, O., Nadler-Yona, C., Nov, S., Koby, S., Baruch, K., Altuvia, S., Elgrably-Weiss, M., Abe, C.M., et al. (2005). Identification of an *Escherichia coli* operon required for formation of the O-antigen capsule. J Bacteriol. 187(15), 5259–5266. 10.1128/JB.187.15.5259-5266.2005.

15. Lu, T., Das, S., Howlader, D.R., Picking, W.D., and Picking, W.L. (2024). *Shigella* Vaccines: The Continuing Unmet Challenge. Int J Mol Sci. 25(8), 4329. 10.3390/ijms25084329.

16. Murayama SY, S.T., Makino S, Kurata T, Sasakawa C, Yoshikawa M. (1986). The use of mice in the Sereny test as a virulence assay of shigellae and enteroinvasive *Escherichia coli*. Infect Immun. 51(2), 696–698. 10.1128/iai.51.2.696-698.1986.

17. van de Verg LL, M.C., Collins HH, Larsen T, Hammack C, Hale TL. (1995). Antibody and cytokine responses in a mouse pulmonary model of *Shigella flexneri* serotype 2a infection. Infect Immun. 63(5), 1947–1954 10.1128/iai.63.5.1947-1954.1995.

18. Shim DH, S.T., Chang SY, et al. (2007). New animal model of shigellosis in the Guinea pig: its usefulness for protective efficacy studies. J Immunol. 178(4), 2476–2482 10.4049/jimmunol.178.4.2476.

19. Miles, S.L., Holt, K.E., and Mostowy, S. (2024). Recent advances in modelling *Shigella* infection. Trends Microbiol. 32(9), 917–924. 10.1016/j.tim.2024.02.004.

20. Alphonse, N., and Odendall, C. (2023). Animal models of shigellosis: a historical overview. Curr Opin Immunol. 85, 102399. 10.1016/j.coi.2023.102399.

21. Ph, Q.S.M., Ledwaba, S.E., Bolick, D.T., Giallourou, N., Yum, L.K., Costa, D.V.S., Oria, R.B., Barry, E.M., Swann, J.R., Lima, A.A.M., et al. (2019). A murine model of diarrhea, growth impairment and metabolic disturbances with *Shigella flexneri* infection and the role of zinc deficiency. Gut Microbes. 10(5), 615–630. 10.1080/19490976.2018.1564430.

22. Howlader DR, B.U., Halder P, Satpathy A, Sarkar S, Ghoshal M, Maiti S, Withey JH, Mitobe J, Dutta S, Koley H. An Experimental Adult Zebrafish Model for *Shigella* Pathogenesis, Transmission, and Vaccine Efficacy Studies. Microbiol Spectr. 10(3), e0034722 10.1128/spectrum.00347-22.

23. Anderson, M.C., Vonaesch, P., Saffarian, A., Marteyn, B.S., and Sansonetti, P.J. (2017). *Shigella sonnei* Encodes a Functional T6SS Used for Interbacterial Competition and Niche Occupancy. Cell Host Microbe. 21(6), 769–776 e763. 10.1016/j.chom.2017.05.004.

24. LaRock, D.L., Chaudhary, A., and Miller, S.I. (2015). *Salmonellae* interactions with host processes. Nat Rev Microbiol. 13(4), 191–205. 10.1038/nrmicro3420.

25. Lorkowski, M., Felipe-Lopez, A., Danzer, C.A., Hansmeier, N., and Hensel, M. (2014). *Salmonella enterica* invasion of polarized epithelial cells is a highly cooperative effort. Infect Immun. 82(6), 2657–2667. 10.1128/IAI.00023-14.

26. Radtke, A.L., Wilson, J.W., Sarker, S., and Nickerson, C.A. (2010). Analysis of interactions of *Salmonella* type three secretion mutants with 3-D intestinal epithelial cells. PLoS One 5(12), e15750. 10.1371/journal.pone.0015750.

27. Caballero-Flores, G., Pickard, J.M., and Nunez, G. (2023). Microbiota-mediated colonization resistance: mechanisms and regulation. Nat Rev Microbiol. 21(6), 347–360. 10.1038/s41579-022-00833-7.

28. Fan, Y., and Pedersen, O. (2021). Gut microbiota in human metabolic health and disease. Nat Rev Microbiol 19(1), 55–71. 10.1038/s41579-020-0433-9.

29. Mullineaux-Sanders, C., Suez, J., Elinav, E., and Frankel, G. (2018). Sieving through gut models of colonization resistance. Nat Microbiol 3(2), 132–140. 10.1128/MMBR.00007-19.

30. Lee, S.M., Donaldson, G.P., Mikulski, Z., Boyajian, S., Ley, K., and Mazmanian, S.K. (2013). Bacterial colonization factors control specificity and stability of the gut microbiota. Nature 501(7467), 426–429. 10.1038/nature12447.

31. Granato, E.T., Smith, W.P.J., and Foster, K.R. (2023). Collective protection against the type VI secretion system in bacteria. ISME J 17(7), 1052–1062. 10.1038/s41396-023-01401-4.

32. Song, L., Pan, J., Yang, Y., Zhang, Z., Cui, R., Jia, S., Wang, Z., Yang, C., Xu, L., Dong, T.G., et al. (2021). Contact-independent killing mediated by a T6SS effector with intrinsic cell-entry properties. Nat Commun. 12(1), 423. 10.1038/s41467-020-20726-8.

33. Tinevez, J.Y., Arena, E.T., Anderson, M., Nigro, G., Injarabian, L., Andre, A., Ferrari, M., Campbell-Valois, F.X., Devin, A., Shorte, S.L., et al. (2019). Tinevez, J.Y., Arena, E.T., Anderson, M., Nigro, G., Injarabian, L., Andre, A., Ferrari, M., Campbell-Valois, F.X., Devin, A., Shorte, S.L., et al. (2019). *Shigella*-mediated oxygen depletion is essential for intestinal mucosa colonization. Nat Microbiol 4(11), 2001–2009. 10.1038/s41564-019-0525-3.

34. Santus, W., Rana, A.P., Devlin, J.R., Kiernan, K.A., Jacob, C.C., Tjokrosurjo, J., Underhill, D.M., and Behnsen, J. (2022). Mycobiota and diet-derived fungal xenosiderophores promote *Salmonella* gastrointestinal colonization. Nat Microbiol 7(12), 2025–2038. 10.1038/s41564-022-01267-w.

35. Kramer, J., Ozkaya, O., and Kummerli, R. (2020). Bacterial siderophores in community and host interactions. Nat Rev Microbiol. 18(3), 152–163. 10.1038/s41579-019-0284-4.

36. Mitchell, P.S., Roncaioli, J.L., Turcotte, E.A., Goers, L., Chavez, R.A., Lee, A.Y., Lesser, C.F., Rauch, I., and Vance, R.E. (2020). NAIP-NLRC4-deficient mice are susceptible to shigellosis. Elife 9, e59022. 10.7554/eLife.59022.

37. Turcotte, E.A., Kim, K., Eislmayr, K.D., Goers, L., Mitchell, P.S., Lesser, C.F., and Vance, R.E. (2026). *Shigella* OspF blocks rapid p38-dependent priming of the NAIP-NLRC4 inflammasome. Proc Natl Acad Sci U S A. 123(3), e2510950123. 10.1101/2025.02.01.636075.

38. Bjursten, M., Hultgren, O.H., and Hultgren Hornquist, E. (2004). Enhanced pro-inflammatory cytokine production in Galphai2-deficient mice on colitis prone and colitis resistant 129Sv genetic backgrounds. Cell Immunol. 228(2), 77–80. 10.1016/j.cellimm.2004.05.001.

39. Kim, S.C., Tonkonogy, S.L., Albright, C.A., Tsang, J., Balish, E.J., Braun, J., Huycke, M.M., and Sartor, R.B. (2005). Variable phenotypes of enterocolitis in interleukin 10-deficient mice monoassociated with two different commensal bacteria. Gastroenterology 128(4), 891–906. 10.1053/j.gastro.2005.02.009.

40. Schmitz, J.M., Tonkonogy, S.L., Dogan, B., Leblond, A., Whitehead, K.J., Kim, S.C., Simpson, K.W., and Sartor, R.B. (2019). Murine Adherent and Invasive *E. coli* Induces Chronic Inflammation and Immune Responses in the Small and Large Intestines of Monoassociated IL-10-/- Mice Independent of Long Polar Fimbriae Adhesin A. Inflamm Bowel Dis. 25(5), 875–885. 10.1093/ibd/izy386.

41. Stefanovic, A., Alam, M.E., Matic, N., Larnder, A., Ritchie, G., Gowland, L., Chorlton, S.D., Lloyd-Smith, E., Payne, M., Dawar, M., et al. (2025). Increased Severity of Multidrug-Resistant *Shigella sonnei* Infections in People Experiencing Homelessness. Clin Infect Dis 80(2), 339–346. 10.1093/cid/ciae575.

42. Lefevre, S., Njamkepo, E., Feldman, S., Ruckly, C., Carle, I., Lejay-Collin, M., Fabre, L., Yassine, I., Frezal, L., Pardos de la Gandara, M., et al. (2023). Rapid emergence of extensively drug-resistant *Shigella sonnei* in France. Nat Commun. 14(1), 462. 10.1038/s41467-023-36222-8.

43. Trivett, H. (2022). Increase in extensively drug resistant *Shigella sonnei* in Europe. Lancet Microbe. 3(7), e481. 10.1016/S2666-5247(22)00160-4.

44. Holt, K.E., Baker, S., Weill, F.X., Holmes, E.C., Kitchen, A., Yu, J., Sangal, V., Brown, D.J., Coia, J.E., Kim, D.W., et al. (2012). *Shigella sonnei* genome sequencing and phylogenetic analysis indicate recent global dissemination from Europe. Nat Genet. 44(9), 1056–1059. 10.1038/ng.2369.

45. Hollingshead, S., McVicker, G., Nielsen, M. R., Zhang, Y., Pilla, G., Jones, R. A., Thomas, J. C., Johansen, S. E. H., Exley, R. M., Brodersen, D. E., & Tang, C. M. (2025). Shared mechanisms of enhanced plasmid maintenance and antibiotic tolerance mediated by the VapBC toxin:antitoxin system. mBio 16(2), e0261624. 10.1128/mbio.02616-24.

46. Sansonetti, P. J., Kopecko, D. J., & Formal, S. B. (1982). Involvement of a plasmid in the invasive ability of *Shigella flexneri*. Infect Immun 35(3), 852–860. 10.1128/iai.35.3.852-860.1982.

47. El-Sayed Ahmed, M.A.E., Zhong, L.L., Shen, C., Yang, Y., Doi, Y., and Tian, G.B. (2020). Colistin and its role in the Era of antibiotic resistance: an extended review (2000-2019). Emerg Microbes Infect 9(1), 868–885. 10.1080/22221751.2020.1754133.

48. Ashida, H., Mimuro, H., and Sasakawa, C. (2015). *Shigella* manipulates host immune responses by delivering effector proteins with specific roles. Front Immunol. 6, 219. 10.3389/fimmu.2015.00219.

49. Moschen, A.R., Adolph, T.E., Gerner, R.R., Wieser, V., and Tilg, H. (2017). Lipocalin-2: A Master Mediator of Intestinal and Metabolic Inflammation. Trends Endocrinol Metab. 28(5), 388–397. 10.1016/j.tem.2017.01.003.

50. Autenrieth, D.M., and Baumgart, D.C. (2012). Toxic megacolon. Inflamm Bowel Dis. 18(3), 584–591. 10.1002/ibd.21847.

51. Clarkson, K.A., Porter, C.K., Talaat, K.R., Frenck, R.W., Jr., Alaimo, C., Martin, P., Bourgeois, A.L., and Kaminski, R.W. (2021). *Shigella*-Specific Immune Profiles Induced after Parenteral Immunization or Oral Challenge with Either *Shigella flexneri* 2a or *Shigella sonnei*. mSphere 6(4), e0012221. 10.1128/mSphere.00122-21.

52. Schroeder, G.N., and Hilbi, H. (2008). Molecular pathogenesis of *Shigella* spp.: controlling host cell signaling, invasion, and death by type III secretion. Clin Microbiol Rev. 21(1), 134–156. 10.1128/CMR.00032-07.

53. Wright, S.S., and Vanaja, S.K. (2022). *Shigella* “Osp”pression of innate immunity. Cell 185(13), 2205–2207. 10.1016/j.cell.2022.05.027.

54. Gamian, A., and Romanowska, E. (1982). The core structure of *Shigella sonnei* lipopolysaccharide and the linkage between O-specific polysaccharide and the core region. Eur J Biochem. 129(1), 105–109. 10.1111/j.1432-1033.1982.tb07027.x.

55. Nadler, C., Koby, S., Peleg, A., Johnson, A.C., Suddala, K.C., Sathiyamoorthy, K., Smith, B.E., Saper, M.A., and Rosenshine, I. (2012). Cycling of Etk and Etp phosphorylation states is involved in formation of group 4 capsule by *Escherichia coli*. PLoS One 7(6), e37984. 10.1371/journal.pone.0037984.

56. Carmen R. Beuzón, D.W.H. (2001). Use of mixed infections with *Salmonella* strains to study virulence genes and their interactions in vivo. Microbes Infect. 3(14-15), 1345–1352. 10.1016/s1286-4579(01)01496-4.

57. Dominguez-Medina, C.C., Perez-Toledo, M., Schager, A.E., Marshall, J.L., Cook, C.N., Bobat, S., Hwang, H., Chun, B.J., Logan, E., Bryant, J.A., et al. (2020). Outer membrane protein size and LPS O-antigen define protective antibody targeting to the *Salmonella* surface. Nat Commun. 11(1), 851. 10.1038/s41467-020-14655-9.

58. Crawford, R.W., Keestra, A.M., Winter, S.E., Xavier, M.N., Tsolis, R.M., Tolstikov, V., and Baumler, A.J. (2012). Very long O-antigen chains enhance fitness during *Salmonella*-induced colitis by increasing bile resistance. PLoS Pathog. 8(9), e1002918. 10.1371/journal.ppat.1002918.

59. Tsai, C.E., Wang, F.Q., Yang, C.W., Yang, L.L., Nguyen, T.V., Chen, Y.C., Chen, P.Y., Hwang, I.S., and Ting, S.Y. (2025). Surface-mediated bacteriophage defense incurs fitness tradeoffs for interbacterial antagonism. EMBO J 44(9), 2473–2500. 10.1038/s44318-025-00406-3.

60. Cota, I., Sanchez-Romero, M.A., Hernandez, S.B., Pucciarelli, M.G., Garcia-Del Portillo, F., and Casadesus, J. (2015). Epigenetic Control of *Salmonella enterica* O-Antigen Chain Length: A Tradeoff between Virulence and Bacteriophage Resistance. PLoS Genet. 11(11), e1005667. 10.1371/journal.pgen.1005667.

61. Ridelfi, M., Vezzani, G., Roscioli, E., Batani, G., Boero, E., Nannini, F., Marini, E., Bhaumik, U., Desalegn, G., Serpino, O., et al. (2025). A multifunctional anti-O-Antigen human monoclonal antibody protects against *Shigella sonnei* infection in vivo. Proc Natl Acad Sci U S A 122(30), e2426211122. 10.1073/pnas.2426211122.

62. Rogawski McQuade, E.T., Liu, J., Mahfuz, M., Havt, A., Varghese, T., Shrestha, J., Kabir, F., Penataro Yori, P., Samie, A., Saidi, Q., et al. (2025). Epidemiology of *Shigella* species and serotypes in children: a retrospective substudy of the MAL-ED observational birth cohort study. Lancet Microbe 6(6), 101064. 10.1016/j.lanmic.2024.101064.

63. Shrum Davis, S., Salazar-Hamm, P., Edge, K., Hanosh, T., Houston, J., Griego-Fisher, A., Lugo, F., Wenzel, N., Malone, D.E., Bradford, C., et al. (2025). Multidrug-resistant *Shigella flexneri* outbreak affecting humans and non-human primates in New Mexico, USA. Nat Commun.16(1), 4680. 10.1038/s41467-025-59766-3.

64. Vacca, M., Celano, G., Calabrese, F.M., Portincasa, P., Gobbetti, M., and De Angelis, M. (2020). The Controversial Role of Human Gut *Lachnospiraceae*. Microorganisms 8(4), 573. 10.3390/microorganisms8040573.

65. Kobayashi, T., Ogawa, M., Sanada, T., Mimuro, H., Kim, M., Ashida, H., Akakura, R., Yoshida, M., Kawalec, M., Reichhart, J.M., et al. (2013). The *Shigella* OspC3 effector inhibits caspase-4, antagonizes inflammatory cell death, and promotes epithelial infection. Cell Host Microbe 13(5), 570–583. 10.1016/j.chom.2013.04.012.

66. Cohen, D., Ashkenazi, S., Green, M.S., Gdalevich, M., Robin, G., Slepon, R., Yavzori, M., Orr, N., Block, C., Ashkenazi, I., et al. (1997). Double-blind vaccine-controlled randomised efficacy trial of an investigational *Shigella sonnei* conjugate vaccine in young adults. Lancet 349(9046), 155–159. 10.1016/S0140-6736(96)06255-1

67. Mancini, F., Gasperini, G., Rossi, O., Aruta, M.G., Raso, M.M., Alfini, R., Biagini, M., Necchi, F., and Micoli, F. (2021). Dissecting the contribution of O-Antigen and proteins to the immunogenicity of *Shigella sonnei* generalized modules for membrane antigens (GMMA). Sci Rep. 11(1), 906. 10.1038/s41598-020-80421-y.

68. Weiserova, M., Dutta, C.F., and Firman, K. (2000). A novel mutant of the type I restriction-modification enzyme EcoR124I is altered at a key stage of the subunit assembly pathway. J Mol Biol. 304(3), 301–310. 10.1006/jmbi.2000.4219.

69. Jackson, S.A., Fellows, B.J., and Fineran, P.C. (2020). Complete Genome Sequences of the *Escherichia coli* Donor Strains ST18 and MFDpir. Microbiol Resour Announc 9(45). 10.1128/MRA.01014-20.

70. Liu, G., Beaton, S.E., Grieve, A.G., Evans, R., Rogers, M., Strisovsky, K., Armstrong, F.A., Freeman, M., Exley, R.M., and Tang, C.M. (2020). Bacterial rhomboid proteases mediate quality control of orphan membrane proteins. EMBO J. 39(10), e102922. 10.15252/embj.2019102922.

71. Bolger, A.M., Lohse, M., and Usadel, B. (2014). Trimmomatic: a flexible trimmer for Illumina sequence data. Bioinformatics 30(15), 2114–2120. 10.1093/bioinformatics/btu170.

72. Ruscheweyh, H.J., Milanese, A., Paoli, L., Karcher, N., Clayssen, Q., Keller, M.I., Wirbel, J., Bork, P., Mende, D.R., Zeller, G., and Sunagawa, S. (2022). Cultivation-independent genomes greatly expand taxonomic-profiling capabilities of mOTUs across various environments. Microbiome 10(1), 212. 10.1186/s40168-022-01410-z.

73. Moor, K., Fadlallah, J., Toska, A., Sterlin, D., Balmer, M.L., Macpherson, A.J., Gorochov, G., Larsen, M., and Slack, E. (2016). Analysis of bacterial-surface-specific antibodies in body fluids using bacterial flow cytometry. Nat Protoc 11(8), 1531–1553. 10.1038/nprot.2016.091. 10.1038/nprot.2016.091.

