## Supplemental figures and tables for "A murine model of *Shigella sonnei* intestinal colonization"

**SUPPLEMENTARY MATERIALS**

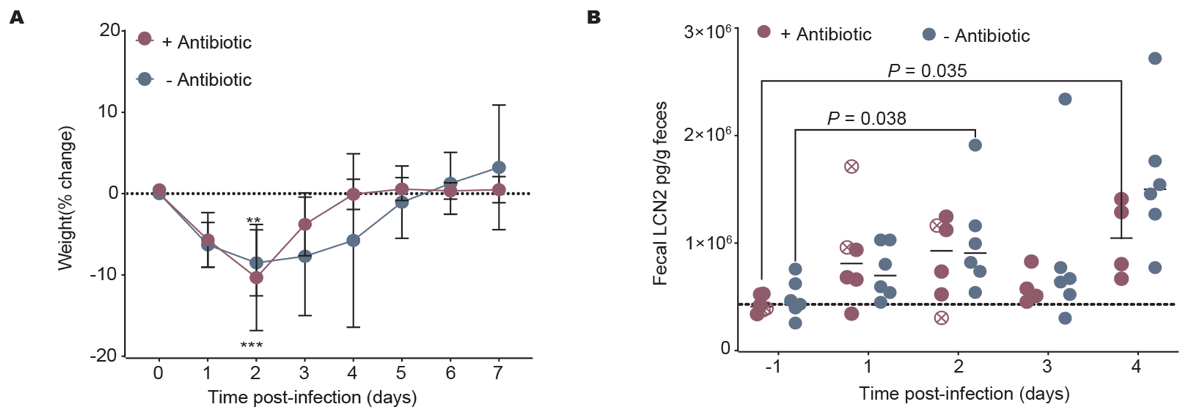

**Figure S1 *Shigella* infection with or without antibiotic pretreatment. A)** Weight of mice infected with *S. sonnei* CS14 with or without antibiotics pretreatment. Dot, mean; error bars, SD. **B)** Fecal LCN2 of infected mice with or without antibiotics pretreatment. Dots, individual mouse; the dot with cross showing significant higher lipocalin at humane endpoint, *n* = 6; Two-way ANOVA, multiple comparisons.

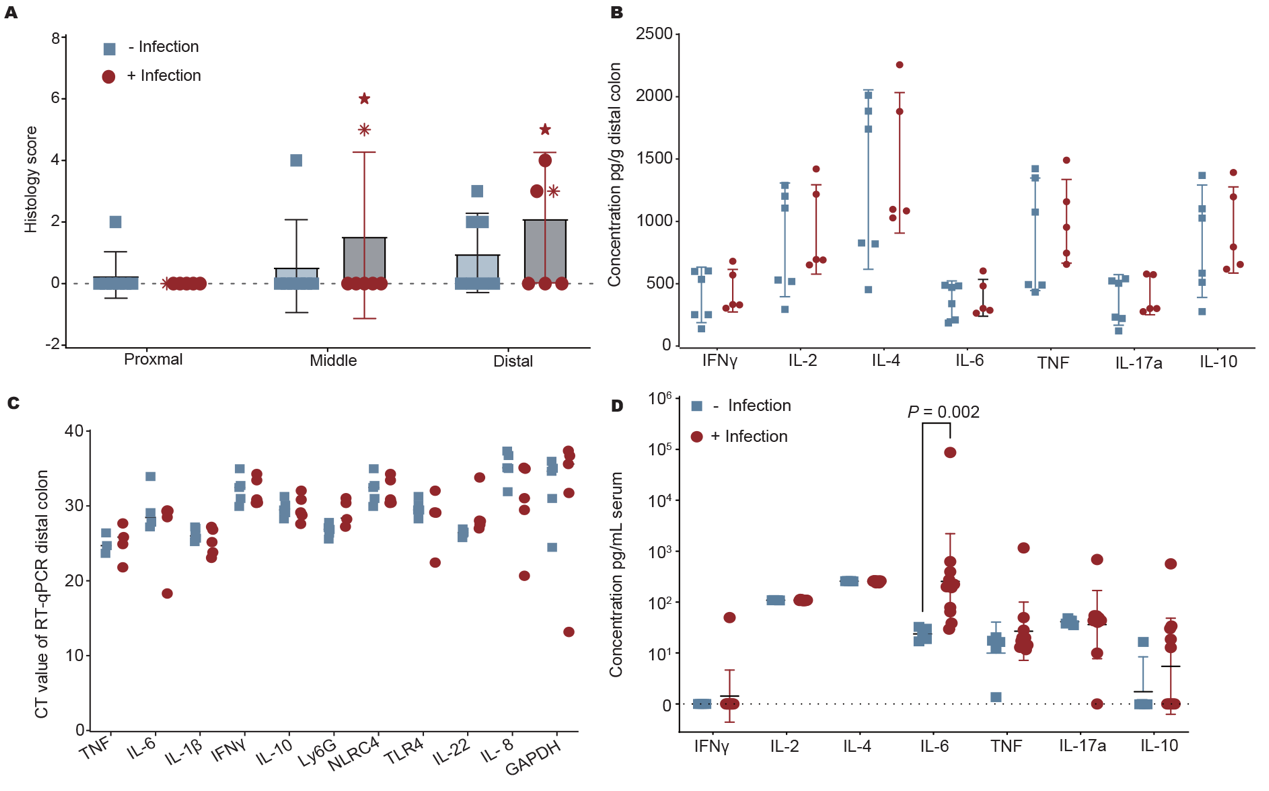

**Figure S2 *Shigella* infection induces individual minor inflammatory response. A)** Histopathology scoring of mice intestinal tissue from the proximal, middle, and distal colon of mice on day 2 uninfected or infected with *S. sonnei*. Each symbol represents an individual mouse; *n* = 7. Cytokine (**B**) and mRNA (**C**) levels in the distal colon or cytokines in the serum (**D**) on day 2 post-infection in *S. sonnei* infected (*n* = 11) and uninfected mice (*n* = 5). Dots, individual mice; Mann-Whitney U.

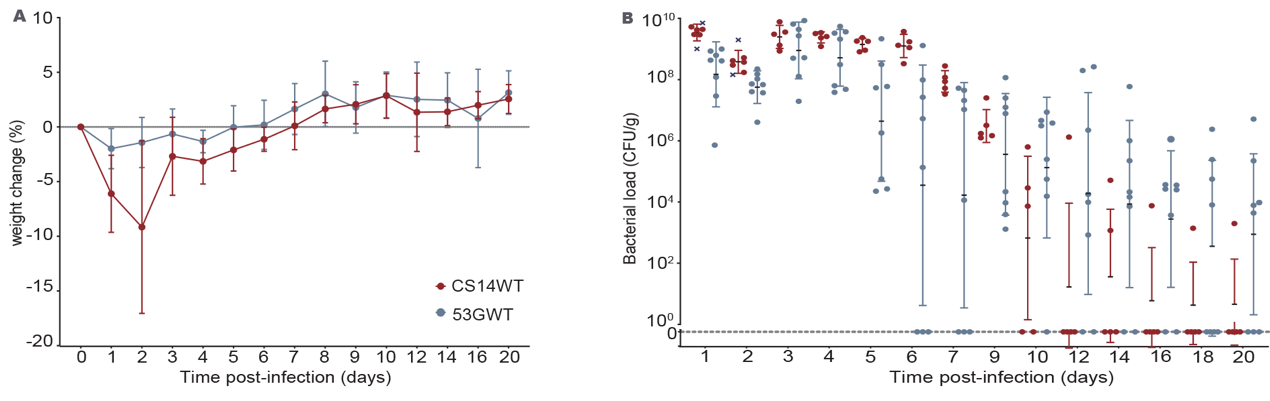
**Figure S3 Challenge mice with different *S. sonnei* strains. A)** Weight of mice infected with *S. sonnei* CS14 or *S. sonnei* 53G (*n* = 7 and 8, respectively); dots, mean; error bars, SD; Mann-Whitney U. **B)** Intestinal colonization with *S. sonnei* CS14 or *S. sonnei* 53G. Dashed line, limit of detection; cross, euthanized mice; dots, individual mice.

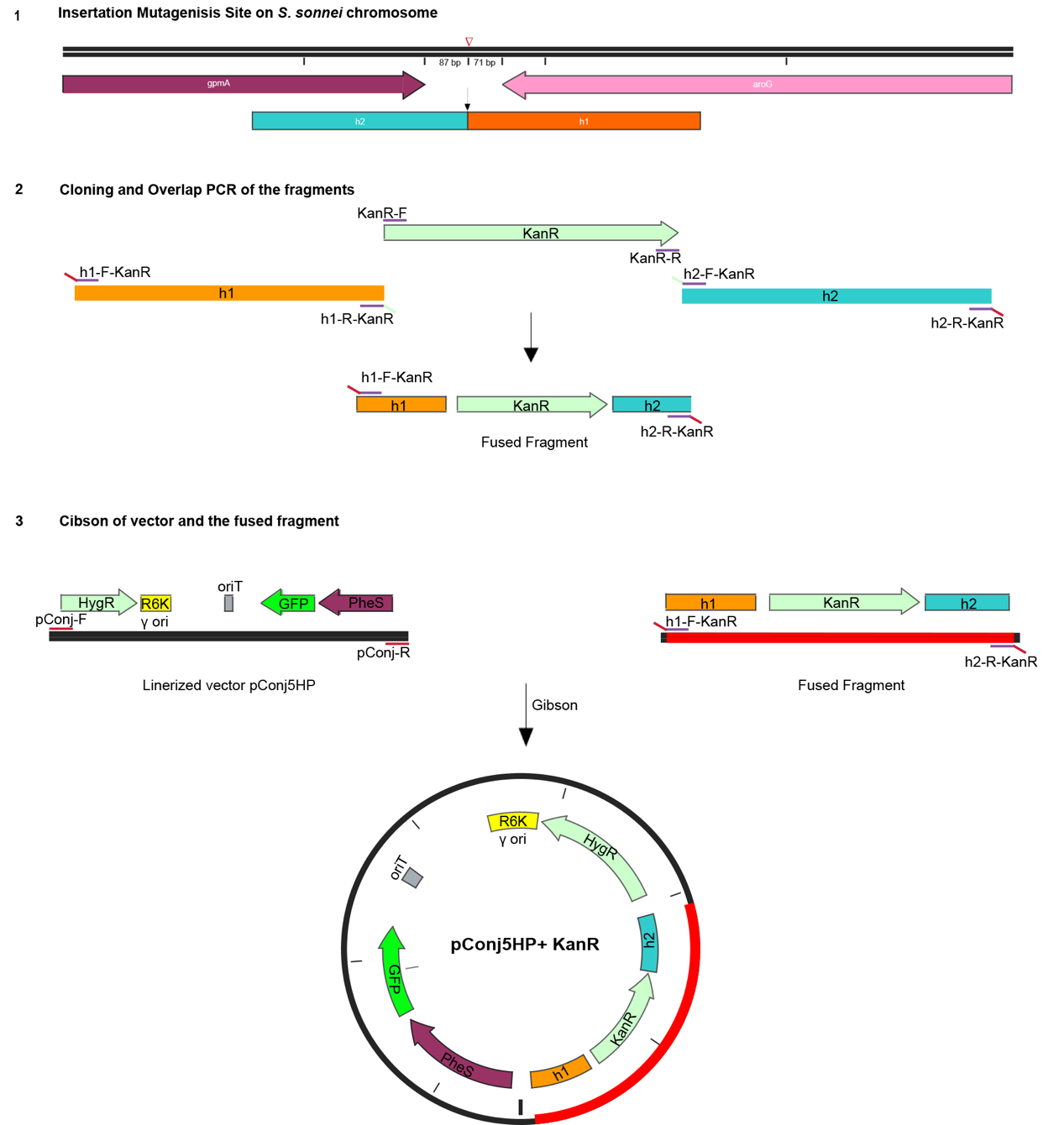

**Figure S4 Introduction of a kan^R^ cassette into the *S. sonnei* CS14 chromosome.** 1. Organisation of *gpmA* and *aroG* which are orientated in a tail-to-tail fashion with a 158 bp intergenic region, and the h1 and h2 fragments amplified. Insertion site, ∇. 2. h1 and h2 from up- and downstream of the insertion site of the kanamycin resistance cassette (Kan^R^) included in overlap PCRs. 3. Gibson assembly of linearized pConj5HP with elements such as Hyg^R^, GFP, and *pheS* (for counter selection) shown.

Table S1 *P* value of the family absolute abundance from day -4 to day 0

| **Main Family** | **Description** | ***P* value** |
| --- | --- | --- |
| Aerococcaceae | + Antibiotic | 1.000 |
|  | - Antibiotic | 1.000 |
| Clostridiaceae | + Antibiotic | 1.000 |
|  | - Antibiotic | 0.562 |
| Coprobacillaceae | + Antibiotic | 0.031 |
|  | - Antibiotic | 0.312 |
| Eggerthellaceae | + Antibiotic | 0.031 |
|  | - Antibiotic | 0.437 |
| Erysipelotrichaceae | + Antibiotic | 1.000 |
|  | - Antibiotic | 0.625 |
| Lachnospiraceae | + Antibiotic | 0.031 |
|  | - Antibiotic | 0.437 |
| Lactobacillaceae | + Antibiotic | 0.062 |
|  | - Antibiotic | 0.312 |
| Muribaculaceae | + Antibiotic | 0.031 |
|  | - Antibiotic | 1.000 |
| Oscillospiraceae | + Antibiotic | 0.031 |
|  | - Antibiotic | 0.437 |
| Peptostreptococcaceae | + Antibiotic | 0.250 |
|  | - Antibiotic | 1.000 |
| Rikenellaceae | + Antibiotic | 0.312 |
|  | - Antibiotic | 0.687 |
| Ruminococcaceae | + Antibiotic | 0.031 |
|  | - Antibiotic | 0.218 |

Table S2 Bacterial strains and plasmids used in this study

| **Strain/Plasmid** | **Description** | **Reference** |
| --- | --- | --- |
| *S. sonnei* CS14 | With Amp^R^ on chromosome | this work |
| *S. sonnei* 53G | With Kan^R^ on chromosome | this work |
| *S. sonnei* OAg*-* | Isolated from HT29 cells |  |
| *S. sonnei* G4C- | Constructed capsule mutant | this work |
| *S. sonnei* T3SS- | Isolated from mice |  |
| *E. coli* | MFD*pir* (∆*hsdR*) | this work |
| pConj5K | Conjugative suicide plasmid with *sacB* and Kan^R^ | this work |
| pConj5HP | Conjugative suicide plasmid with *sacB* and Hyg^R^ | this work |

Table S3 Primers used in this study

| **Primer name** | **Sequence (5' - 3')** | **Purpose** |
| --- | --- | --- |
| pConj-F | AAACGTCATAGCTGTTTCCTGCTAGTATAG | Plasmid linearization |
| pConj-R | AAACACTGGCCGTCGTTTTACTATACTCC |  |
| aroG-h1F-Amp | CGGAGTATAGTAAAACGACGGCCAGTGTTTTAACTAAATGGGGGCATTCGG | Cloning arog fragments for plasmid and chromosome homology sequence recombination |
| aroG-h1R-Amp | TTAGAAAAATAAACAAATAGGGGTTCCGCGCTGCGCGTCTTATCAGGC |  |
| aroG-h2F-Amp | CGCGGAACCCCTATTTGTTTATTTTTC |  |
| aroG-h2R-Amp | TTACCAATGCTTAATCAGTGAGGC |  |
| seq-F | CACTAAATAATAGTGAACGGCAGG | Plasmid sequencing |
| seq-R | CACTGATGAGAATATCGTCGG |  |
| Amp^R^-F | CGCGGAACCCCTATTTGTTTATTTTTC | Cloning amp sequencing for insertion to chromosome |
| Amp^R^-R | TTACCAATGCTTAATCAGTGAGGC |  |
| KanR-F | TTCAAATATGTATCCGCTCATGAGACAATAAC | Cloning Km sequencing for insertion to chromosome |
| KanR-R | ATCGGGCCGGATCTAGATATTTAG |  |
| h1F-KanR | CGGAGTATAGTAAAACGACGGCCAGTGTTTGGGGGCATTCGGCGATTG | Cloning arog flank fragments for plasmid and chromosome homology sequence recombination |
| h1R-KanR | GGGTTATTGTCTCATGAGCGGATACATATTTGAACTGCGCGTCTTATCAGGCC |  |
| h2F-KanR | AGTTTTTCTAAATATCTAGATCCGGCCCGATCGTCGCATCAGGCAATGTG |  |
| h2R-KanR | CTATACTAGCAGGAAACAGCTATGACGTTTGTTATCCGGGTCACGATCCGC |  |
| g4c-h1F | CGGAGTATAGTAAAACGACGGCCAGTGTTTGCTTTATTTTATTAACCCCGTATAGTGCAG | Cloning g4c flank fragments for plasmid and chromosome homology sequence recombination |
| g4c-h1R | TGGCGTCGTCAACATGAAGG ∇ |  |
| g4c-h2F | CATGTTGACGACGCCACGTCAATATTAAAGGCGCTATCCTC |  |
| g4c-h2R | CTATACTAGCAGGAAACAGCTATGACGTTTGAGATATTTCTCCTGCAACAAGCAC |  |
| g4c-seqF | GGCGGTCTTAAATATCGCTGG | sequencing for g4c mutant |
| g4c-seqR | GCCGACAATGTCTCATTAACCG |  |
| TNF-α-F | CCCTCACACTCAGATCATCTTCT | RT-qPCR to detect mice cytokine expression |
| TNF-α-R | GCTACGACGTGGGCTACAG |  |
| IL-6-F | CCGGAGAGGAGACTTCACAG |  |
| IL-6-R | TCCACGATTTCCCAGAGAAC |  |
| IL-1β-F | GCAACTGTTCCTGAACTCAACT |  |
| IL-1β-R | ATCTTTTGGGGTCCGTCAACT |  |
| IFN-γ-F | TCAAGTGGCATAGATGTGGAAGAA |  |
| IFN-γ-R | TGGCTCTGCAGGATTTTCATG |  |
| IL-10-F | GCTCTTACTGACTGGCATGAG |  |
| IL-10-R | CGCAGCTCTAGGAGCATGTG |  |
| IL-8-F | GCTTCTCCTTCTCCACAACC |  |
| IL-8-R | TGAAGGCAAGCCTCGTGGTA |  |
| IL-22-F | TGTGCCAGGAGGGTGCTGTT |  |
| IL-22-R | GTCGTCGTCTCAAAGTCCAG |  |
| TLR4-F | AGCATTTCAGAGCCGTTGGT |  |
| TLR4-R | CAGGTCCAGGTTCTTGGTTG |  |
| virF_F | CTTAGCTTGTTGCACAGAGA |  |
| vitF-R | AAGATGGGCTTGATATTCCG |  |
| hns-F | GCTCAACAGTATGCACAGAA |  |
| hns-R | TTGCAAAGGCGTTGAATTA |  |
| wzy_F | GGTTTCACGTTTCTCTGTGG |  |
| wzy_R | CCTTTACCAATATACCCTCCGC |  |

Table S4 Histology score system

| Basis for proposed scoring system: | | |  |
| --- | --- | --- | --- |
| A. | EPITHELIUM | HYPERPLASIA | and/or GOBLET CELL DEPLETION |
|  | 0 | None | None |
|  | 1 | Mild (1.5x) | Mild (25%) |
|  | 2 | Moderate (2-3x) | Marked (25-50%) |
|  | 3 | Severe (>3x) | Substantial (>50%) |
| B. | INFLAMMATION IN LAMINA PROPRIA | | |
|  | 0 | None - few leucocytes | |
|  | 1 | Mild - some increase in leucos at tips of crypts OR many lymphoid follicles | |
|  | 2 | Moderate- marked infiltrate (notable broadening of crypt) | |
|  | 3 | Severe -dense infiltrate throughout | |
| C. | AREA AFFECTED (% of section) | | |
|  | 0 | None |  |
|  | 1 | up to 25% |  |
|  | 2 | 25-50% |  |
|  | 3 | >50% |  |
| D. | MARKERS OF SEVERE INFLAMMATION | | |
|  | 0 | None |  |
|  | 1 | Submucosal inflammation OR Few crypt abscesses (<5) | |
|  | 2 | Submucosal inflammation AND Few Crypt Absecesses (<5) | |
|  | 2 | Many crypt abscesses (<5) OR Extensive submucosal inflamm OR Crypt branching (*Hh)* | |
|  | 3 | Many crypt abscesses (<5) AND Extensive submucosal inflamm / Crypt branching (*Hh)* | |
|  | 3 | Ulceration OR Extensive fibrosis | |
